# Exploration of the autonomous replication region and its utilization for expression vectors in cyanobacteria

**DOI:** 10.1101/2022.11.17.516977

**Authors:** Yutaka Sakamaki, Kaisei Maeda, Kaori Nimura-Matsune, Taku Chibazakura, Satoru Watanabe

## Abstract

Due to their photosynthetic capabilities, cyanobacteria is expected to be an ecologically friendly host for the production of biomaterials. However, compared to other bacteria, there is little information of autonomous replication sequences, and tools for genetic engineering, especially expression vector systems, are limited. In this study, we established an effective screening method, namely AR-seq (Autonomous Replication sequencing), for finding autonomous replication regions in cyanobacteria and utilized the region for constructing expression vector. AR-seq using the genomic library of *Synechocystis* sp. PCC 6803 revealed that a certain region containing Rep-related protein (here named as Cyanobacterial Rep protein A2: CyRepA2) exhibits high autonomous replication activity in a heterologous host cyanobacterium, *Synechococcus elongatus* PCC 7942. The reporter assay using GFP showed that the expression vector pYS carrying CyRepA2 can be maintained in a wide range of multiple cyanobacterial species, not only *S*. 6803 and *S*. 7942, but also *Synechococcus* sp. PCC 7002 and *Anabaena* sp. PCC 7120. In *S*. 7942, the GFP expression in pYS-based system can be tightly regulated by IPTG, achieving 10-fold higher levels than that of chromosome-based system. Furthermore, pYS can be used together with conventional vector pEX, which was constructed from an endogenous plasmid in *5*. 7942. The combination of pYS with other vectors is useful for genetic engineering, such as modifying metabolic pathways, and is expected to improve the performance of cyanobacteria as bioproduction chassis.

## 1 Introduction

Cyanobacteria are the predominant phototrophs in ocean and freshwater ecosystems and are one of the most widespread phylogenetic clades. Cyanobacteria have oxygen-producing photosynthetic capabilities, meaning that they can produce biomass by using solar energy, and they have recently gained attention for their potential as green cell factories for CO2-neutral biosynthesis of various products (Knoot et al., 2018) (Farrokh et al., 2019). Recently, three cyanobacterial model strains, *Synechocystis* sp. PCC 6803 (*S*. 6803), *Synechococcus elongatus* PCC 7942 (*S*. 7942) and *Synechococcus* sp. PCC 7002 (*S*. 7002) have been used in synthetic biology studies for biosynthesis of multiple products including free fatty acids (Liu et al., 2011), isoprene (Lindberg et al., 2010), 2,3-butanediol (Oliver et al., 2013), 1-butanol (Lan and Liao, 2011), isobutyraldehyde (Atsumi et al., 2009) and squalene (Englund et al., 2014). The multicellular filamentous cyanobacterium *Anabaena* sp. PCC 7120 (*A*. 7120), performing both nitrogen and carbon fixation, is particularly suitable for the production of nitrogenous substances and has recently been studied for ammonia production (Higo et al., 2016).

In several model cyanobacteria, *S*. 7942, *S*. 6803, and *S*. 7002, integration of genetic construct to chromosomal neutral sites has been used for exogenous gene expression (Watanabe et al., 2012) (Englund et al., 2015) (Chin et al., 2018) (Ishizuka et al., 2006). Due to their polyploidy, chromosomal expression systems in these cyanobacteria are expected to be more effective than those of monoploid organisms, while genetic manipulation is limited to a few cyanobacteria that have abilities of natural competence and chromosomal recombination. In addition, genetic engineering relying on the integration of the neutral sites is time-consuming when performed sequentially, because all chromosomes must be segregated in order to stably retain the genetically modified constructs in polyploid cyanobacteria.

Self-replication plasmids are maintained autonomously using replication initiation factors (e.g. Rep proteins) encoded in the plasmids themselves and the host’s replication machinery. The autonomous replication sequence consists of an initiation factor and its binding sequence, that serves as the replication origin, but the binding sequence can also function alone if the host’s replication initiation factor functions in *trans*. Most plasmid replicons in *E. coli* cannot be used directly in cyanobacteria and there is little information on autonomous replication sequence as well as available plasmid vectors in cyanobacteria. The broad-host-range plasmid RSF1010, a member of the IncQ plasmids, stably replicates in a wide variety of gram-negative and gram-positive bacteria, including *E. coli* and several strains of cyanobacteria (Trieu-Cuot et al., 1987) (Meyer, 2009) (Cassier-Chauvat et al., 2021). The RSF1010 vectors are maintained in *S*. 6803 cells, but not suitable for gene overexpression because their copy number is comparable to that of chromosomes (Jin et al., 2018). Other vectors utilizing cyanobacterial endogenous plasmids have also been developed: pUH24/pANS (the derivative vector named as pUC303) in *S*. 7942 (Kuhlemeier et al., 1983) (Van der Plas et al., 1992), pCA2.4, pCB2.4 (pSOMA series) (Opel et al., 2022) and pCC5.2 (pSCB) (Jin et al., 2018) in *S*. 6803, pAQ (pAQ1-EX1) in *S*. 7002 (Miyasaka et al., 1998) and pDU1 (pRL series) in *Nostoc* sp. PCC 7524 (Wolk et al., 1984). However, these vectors have not been tested for the host range with the exception of the pSOMA, which has been demonstrated recently to be maintained in two cyanobacteria *S*. 6803 and *A*. 7120 (Opel et al., 2022). Further development of cyanobacterial genetic engineering will require more information on the autonomous replication regions that function in a wide range of species.

In this study, we established an effective screening method, namely AR-seq (Autonomous Replication sequencing), for finding autonomous replication regions in cyanobacteria and found that a region containing Rep-related protein ORF B (here named as Cyanobacterial Rep protein A2: CyRepA2) encoded in the plasmid pCC5.2 derived from *S*. 6803 has replication activity in *S*. 7942. Using this region, we constructed an expression vector pYS and tested its expression efficiency, host-range, and compatibility with other plasmids. Our results suggest that they contribute to not only elucidating Rep proteins and their regulatory mechanisms, which are poorly known in cyanobacteria, but also improving the performance of cyanobacteria as bioproduction chassis.

## 2 Material and Methods

### 2.1 Cyanobacteria strains and growth condition

The freshwater cyanobacteria *Synechococcus elongatus* PCC 7942 (our laboratory strain *S*. 7942 TUA, which lacks endogenous small plasmid pHU24/pANS) (Watanabe et al., 2012) and *Synechocystis* sp. PCC 6803 (sub-strain *S*. 6803 GT-I) (Kanesaki et al., 2012) were used in this study. Both strains were cultured in modified BG-11 medium, which contains double amount of sodium nitrate and 20 mM HEPES-KOH (pH 8.2), while *Anabaena* sp. PCC 7120 was maintained in the standard BG11 (Castenholz, 1988). The marine cyanobacterium *Synechococcus* sp. PCC 7002 was cultured in modified A2 medium, which contains the one-third amount of sodium nitrate (Hasunuma et al., 2019). Cultures were grown photoautorophically at 30 °C, which was illuminated continuously (50 μmol photons m^-2^s^-1^) with 2% CO_2_ (v/v) bubbling. When appropriate, spectinomycin (*S*. 7942: final concentration 40 μg/mL, *A*. 7120: 10 μg/mL) or chloramphenicol (*S*. 7942 and *S*. 6803: 7.5 μg/mL, *S*. 7002: 20 μg/mL) were added to medium.

### 2.2 Autonomous replication sequencing (AR-seq)

Each DNA fragments were amplified using PrimeSTAR DNA polymerase (TaKaRa, Shiga, Japan) and subcloned into vectors using T4 DNA ligase (TakaRa) or In-Fusion HD Cloning Kit (TaKaRa). For the construction of a *S*. 6803 genomic library, ColE1 origin and chloramphenicol acetyltransferase gene (*cat*) were PCR-amplified from pUC4K (Amersham, Chicago, IL, USA) and pUC303 (Kuhlemeier et al., 1983) plasmids using respective primer sets (F1/ R2, F3/ R4, Supplementary Table S1) and recombined by PCR (primer, F3/ R2). After digestion by *BamHI* and dephosphorylation treatment using SAP (TaKaRa), the DNA fragment was ligated with 1.5-2.5 kbp fragments of the *Sau*3AI-digested *S*. 6803 genomic DNA. As a result of the transformation of *E. coli*, we obtained 2.5 × 10^4^ clones (Library A). In an attempt to determine the regions responsible for replication in heterologous cyanobacteria, we transformed *S*. 7942 cells with the *S*. 6803 genomic library to yield 335 colonies and pooled them (Library B). To obtain plasmids that replicated independently of chromosomal integration, the DNAs extracted from library B was introduced into *E. coli* again, and 1.4 × 10^4^ transformants were obtained (Library C). The genomic libraries (Libraries A-C) were subjected to comprehensive sequencing analysis.

The sequencing library was constructed by NEB Next Ultra DNA Library Prep Kit (NEB, Ipswich, MA, USA) and analyzed by the MiSeq sequencer using a paired-end 150 bp sequence read run with the MiSeq reagent kit v3 (illumina, San Diego, CA, USA). The numbers of raw read pairs per sample were shown in Supplementary Table S2. The reads were trimmed using CLC Genomics Workbench ver. 9.5.4 (QIAGEN, Venlo, Netherland) with the following parameters: Phred quality score >30; ambiguous nucleotides allowed: 1; automatic read-through adaptor trimming: yes; removing the terminal 15 nucleotides from the 5’ end and 5 nucleotides from the 3’end; and removing truncated reads of less than 30 nucleotides in length. To approximately identify the regions included in the library, trimmed reads were mapped to the *S*. 6803 genome (Accession numbers, chromosome: AP012276, pSYSM: AP004310, pSYSX: AP006585, pSYSA: AP004311, pSYSG: AP004312, pCA2.4: CP003270, pCB2.4: CP003271, and pCC5.2: CP003272) using CLC Genomics Workbench ver. 20.0.1 (QIAGEN) with the following parameters: match score: 1; mismatch cost: 2; indel cost: 3; length fraction: 0.8; similarity fraction: 0.9; and non-specific match handling: ignore. Original sequence reads were deposited in the DRA/SRA database with the following accession numbers (Library A: DRR285589, Library B: DRR285594, and Library C: 285596). The accession number of BioProject was PRJDB11466.

### 2.3 Phylogenetic analysis

The amino acid sequence of ORF B (Accession number: WP_015390179.1) was obtained from the database, and its homologs were retrieved by NCBI-BLAST search for the top 100 most similar sequences. Among them, we excluded the overlapping sequences and obtained 58 homologous sequences. A phylogenetic analysis was performed on these homologs, along with pUH24/pANS Rep in *S*. 7942 and RSF1010 RepC, using ClustalW within MEGA11 with default parameters (Tamura et al., 2021). The evolutionary history was inferred using the Neighbor-Joining method (Saitou and Nei, 1987). The optimal tree is shown in Figure 2. The percentage of replicate trees in which the associated taxa clustered together in the bootstrap test (1000 replicates) are shown next to the branches (Felsenstein, 1985). The tree is drawn to scale, with branch lengths in the same units as those of the evolutionary distances used to infer the phylogenetic tree. The evolutionary distances were computed using the Poisson correction method (Zuckerkandl and Pauling, 1965) and are in the units of the number of amino acid substitutions per site. This analysis involved total 61 amino acid sequences (ORFB, pUH24/pANS Rep, RSF1010 RepC, and 58 homologs). All ambiguous positions were removed for each sequence pair (pairwise deletion option). There were a total of 1655 positions in the final dataset. Evolutionary analyses were conducted in MEGA11 (Tamura et al., 2021).

**Figure 1.**
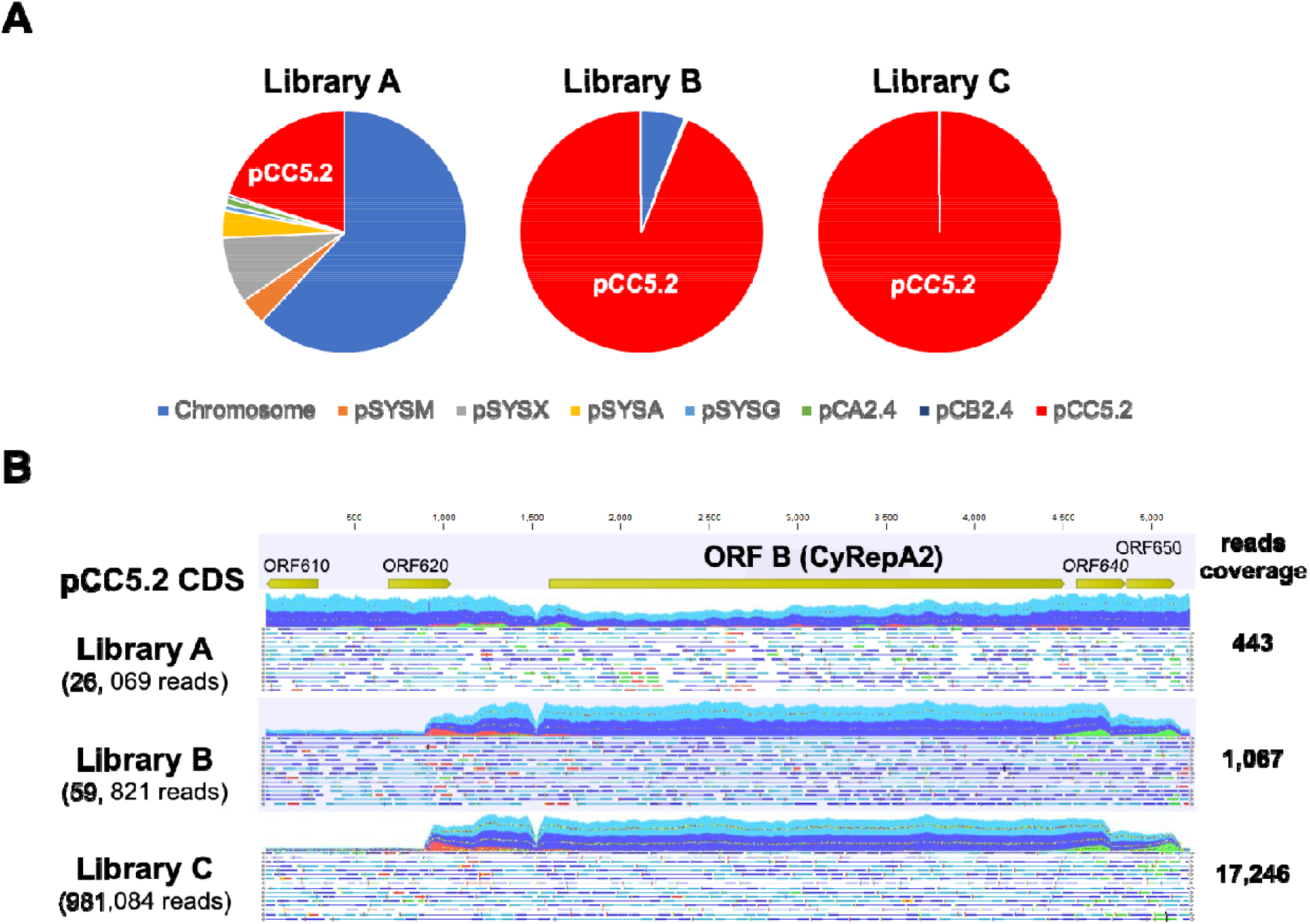
Screening of autonomous replication region in *S*. 7942 cell. (A) The composition of sequence reads of the *S*. 6803 genomic libraries obtained from AR-seq analysis. Three libraries were sequenced by MiSeq and the reads were mapped to the *S*. 6803 genome (chromosome and 7 plasmids, pSYSM, pSYSX, pSYSA, pSYSG, pCA2.4, pCB2.4, and pCC5.2). The sequence reads were analyzed by the CLC Genomics Workbench ver. 20.0.1. Library A: *S*. 6803 genomic library before the screening, Library B: DNA extracted from *S*. 7942 cells transformed Library B, Library C: DNA extracted from *E. coli* cells transformed Library B. (B) Mapping results of sequence reads to pCC5.2 as references. Reads that have been successfully mapped to pCC5.2 as a pair are shown as a blue or light blue, while only one of the reads in the pair has been mapped is shown as a red or green. The reads coverage shows the maximum depth of the mapping reads.

**Figure 2.**
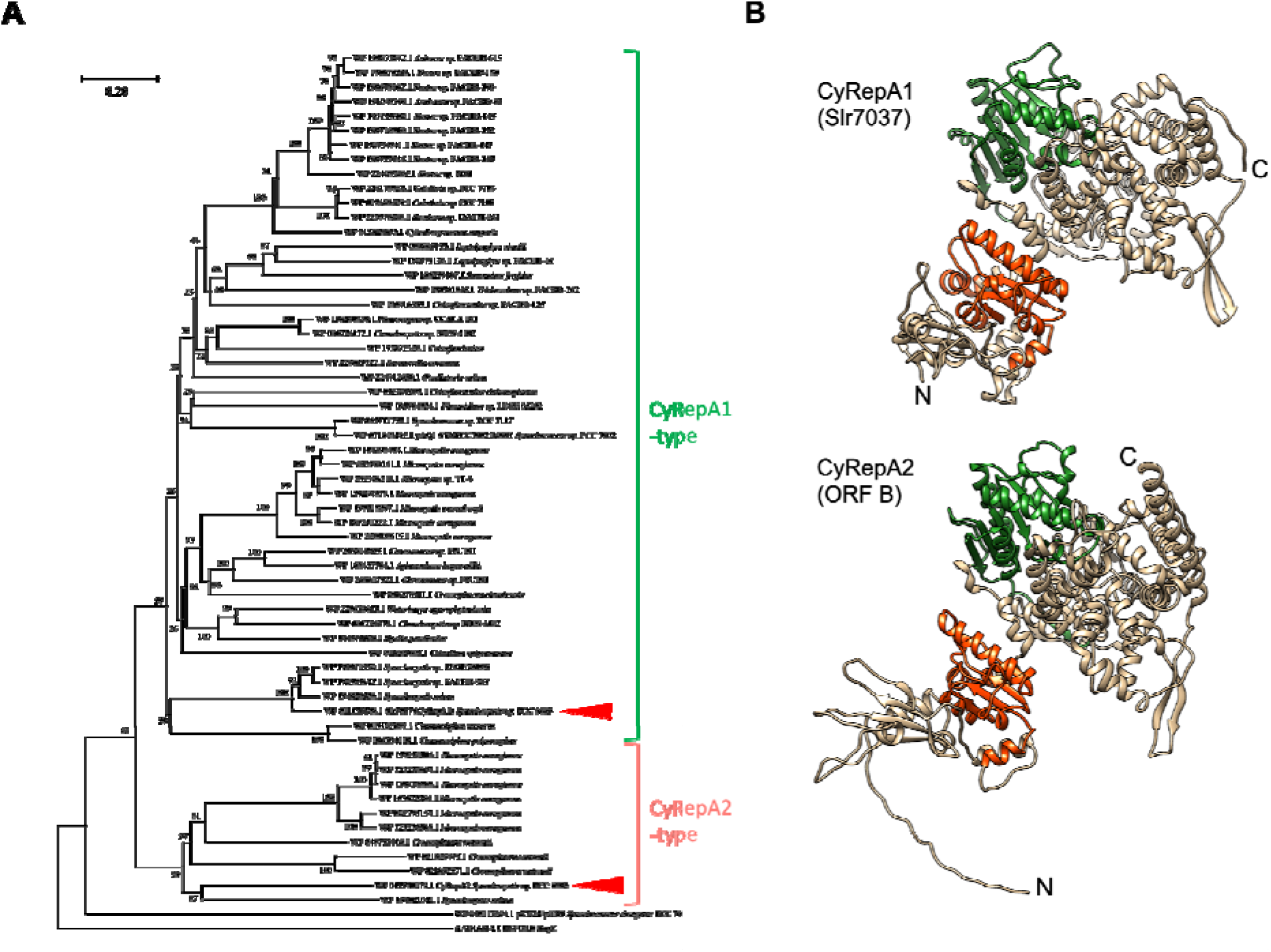
Phylogenetic and structural analysis of CyRepA1 (Slr7037) and CyRep2 (ORF B) (A) Phylogenetic tree of CyRepA. CyRepA2 (ORF B) and its 58 homologues were used phylogenetic analysis along with pUH24/pANS Rep and RSF1010 RepC used as outgroups. Proteins used for analysis were shown by its Accession number in database with organism name. The tree is drawn to scale, with branch lengths in the same units as those of the evolutionary distances used to infer the phylogenetic tree. CyRepA1 (Slr7037) and CyRepA2 are indicated by red triangles. (B) 3D structures of CyRepA1 and CyRepA2 predicted using AlphaFold2. The 3D structures are color-coded as follows based on the results of domain predictions by SMART and Pfam. DUF3854 domain (orange), DEXDc domain (green).

### 2.4 Prediction of 3D structure of CyRepAs

Alphafold2 was used for 3D structure prediction. (AlQuraishi, 2019) (Cramer, 2021). The predicted 3D structure of Slr7037 (CyRepA1) was obtained from the AlphaFold Protein Structure Database. ORF B (CyRepA2) was predicted by ColabFold, and the model with the best score was adopted (Mirdita et al., 2022). The Predicted 3D structures were colored to domains using UCSF Chimera (Pettersen et al., 2004). Domain predictions were performed using SMART and Pfam (Letunic et al., 2021) (Mistry et al., 2021).

### 2.5 Plasmid and strain construction

To construct the expression vector pYS1C-GFP, we used the plasmid isolated from the *E. coli* transformant pools of Library B (Supplementary Table S3). DNA fragments containing *lacI* gene*, trc* promoter, *GFP^mut2^* gene and B0011 *lux* operon terminator were PCR-amplified by appropriate primer sets (F5/ R6, F7/ R8, F9/ R10, F11/ R12, F13/R14) and introduced into the plasmid obtained from library screening and containing ColE1, *cat*, and a part of pCC5.2 (1174-4674 nt) by In-Fusion cloning (Supplementary Figure S1).

In order to strictly control gene expression in *S*. 7002, the promoter region of pYS1C-GFP was replaced with *clac143* promoter, optimized for regulation in *S*. 7002. Using primers F15/ R16 flanking the sequence of *clac143* promoter, DNA fragment was PCR-amplified from pYS1C-GFP and circularized by In-Fusion cloning. The resulting plasmid was named as pYS4C-GFP. To replace the *cat* gene in pYS1C-GFP and pYS4C-GFP to spectinomycin-resistance (*Sp^R^*) gene, the plasmid regions and *Sp^R^* gene were PCR-amplified using appropriate primer sets (F17/ R18, F19 / R20) and combined using In-Fusion cloning. The resulting plasmids were named as pYS1S-GFP and pYS4S-GFP, respectively.

For the construction of integration plasmid pNSG, DNA fragment of *GFP^mut2^* gene (Cormack et al., 1996) was PCR-amplified with primers F21/ R22 and cloned into the *SalI* and *HindIII* site of integration vector pNSE1 (Kato et al., 2008), containing the homologous region of neutral site in *S*. 7942. To construct a plasmid harboring pUC303-based replication system, the region containing *repA* and *repB* of pUC303, p15A origin, and DNA fragment containing *Sp^R^* gene, and *mScarlet* gene under the *cpcB* promoter were PCR-amplified using appropriate primer sets (F23/R24, F25/R26, F27/R28) and combined using In-Fusion cloning (named pEX2S-mScarlet).

To transform the cyanobacteria *S*. 7942, *S*. 6803 and *S*. 7002, cells were grown as described above until they reached OD750 of 0.7-1.2 and then collected by centrifugation at 3000 × *g* for 10 min at 25 °C. After resuspension in BG11 to a 10-fold concentration, plasmid DNA was added and incubated at 25 °C overnight in light-shield and rotating conditions. The samples were then light irradiated for 1 hour and spread on plates containing drug. In the case of *A*. 7120 transformation, the plasmids pYS1S-GFP or pYS4S-GFP were transformed into *E. coli* HB101 carrying the plasmid pRL623 (Elhai et al., 1997) and were transferred to *A*. 7120 cell with the help of the conjugative plasmid RP4 according to the triparental mating method (Elhai and Wolk, 1988) (Thiel and Wolk, 1987) with minor modification. The *E. coli* cultures carrying the donor plasmid were mixed with the *E. coli* culture carrying RP4 and incubated for 1 hour. The *A*. 7120 culture prepared as described above was added to the *E. coli* mixture and incubated overnight on the BG11 plate containing 5% LB medium. The mixture was harvested by adding BG11 medium onto the plate and spread on the plates containing spectinomycin.

The structures of pYS plasmids introduced into cyanobacteria were confirmed by following procedure: DNAs extracted from cyanobacteria harboring pYS plasmids were used to transform *E. coli* JM109. After the selection in the LB medium containing chloramphenicol or spectinomycin, the plasmid DNAs were extracted by using FavorPrep^TM^ Plasmid DNA Extraction Mini Kit (FAVORGEN, Ping Tung, Taiwan). Plasmid DNAs were digested with *NdeI* and *PstI* for pYS1C-GFP and pYS4C-GFP, and *SalI* and *NheI* for pEX2S-mScarlet. 80 ng of DNA was subjected to 1 % agarose gel electrophoresis (Figure 3B and Supplementary Figure S2).

**Figure 3.**
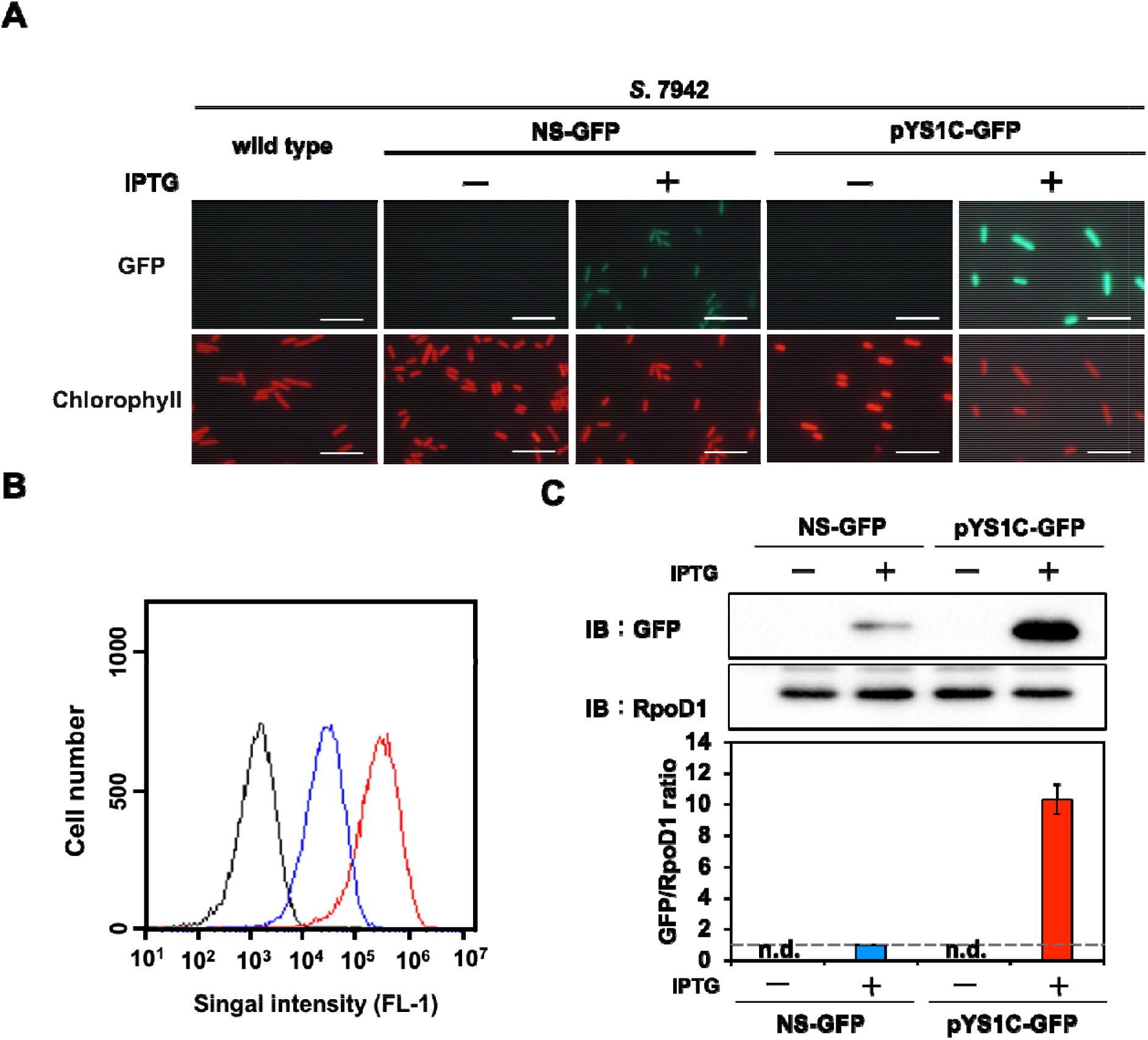
Comparison of expression performance between chromosome and pYS1 plasmid by the GFP reporter system in *S*. 7942. The GFP expression levels in pYS1C-GFP transformant were compared to *S*. 7942 wild-type and NS-GFP strain expressing GFP at chromosomal neutral site. Final 1 mM IPTG was added to induce GFP expression. (A) Fluorescence microscopy. Images of GFP and chlorophyll fluorescence were shown. White bar, 10 μm. (B) FACS analysis of GFP fluorescence. Signal intensity of FL-1 indicating GFP fluorescence in the wild type (black), NS-GFP (blue) and pYS1-GFP (red) strains were shown. (C) Western blotting analysis. The protein extracts obtained from wild type, NS-GFP, and pYS1C-GFP strains were subjected to SDS-PAGE and analyzed by the western blotting using antibodies against GFP and RpoD1 used as an internal control. The signal intensities of GFP were normalized with those of RpoD1, and the ratio of GFP signal of NS-GFP was set to 1. Bars represent mean ± SEM (*n* = 3).

### 2.6 SDS-PAGE and western blotting analysis

The protein extracts from cyanobacteria cells were prepared according to the method described by Nimura et al. (2001) (Nimura et al., 2001) and were separated by SDS-PAGE with 12% Polyacrylamide gel at 150 V for 90-100 min in electrophoresis buffer (25 mM Tris, 192 mM Glycine, 0.1% SDS). The gels were blotted onto 0.2 μm PVDF membranes (Bio-Rad, Hercules, CA, USA) at 2.5 A for 10 min using a Trans-Blot Turbo Transfer System (Bio-Rad). Membranes were blocked for 30 min with 5% skimmed milk in TBS-T (50 mM Tris-HCl (pH 7.5), 150 mM NaCl, 0.05% Tween 20), washed, and soaked in TBS-t for 2 h at room temperature with anti-GFP (MBL, Tokyo, Japan), anti-RpoD1 rabbit antiserum (Seki et al., 2007) or anti-RbcL (Agrisera, Vännäs, Sweden) as primary antibodies. After washing the membranes, they were soaked in TBS-t for 30 min at room temperature with HRP-conjugated anti-mouse (GE Healthcare, Chicago, IL, USA) or anti-rabbit antibody (GE Healthcare) as secondary antibodies. Chemiluminescence was detected with LumiGLO Chemiluminescent Substrate KPL using ChemiDoc XRS Plus (Bio-Rad).

### 2.7 Flowcytometry

Cells were analyzed by GFP fluorescence activated cell sorting (FACS) using BD Accuri™ C6 Flow cytometer (BD Biosciences, San Jose, CA, USA) and the software BD CFlow (BD Biosciences), as described by Watanabe et al. (2012) (Watanabe et al., 2012). Cyanobacterial cells showing chlorophyll fluorescence were sorted by the FL3 channel, and GFP fluorescence was measured by the FL1 channel.

### 2.8 Microscopy

Fluorescent images were obtained using BX53 microscope (OLYMPUS, Tokyo, Japan) at × 40 or 100 magnifications with a DP71 digital camera (OLYMPUS) and DP Controller software ver. 3. 3. 1. 292 (OLYMPUS).

## 3 Results

### 3.1 Autonomous replication sequencing (AR-seq): Library screening of sequences responsible for autonomous replication in cyanobacterial cell

Because of the great diversity of Rep proteins and their regulatory mechanisms, it is impossible to predict a DNA region necessary for autonomous replication based on its sequence alone. To gain new insights into the autonomous replication region in cyanobacteria, we established the new method, namely AR-seq (<u>A</u>utonomous <u>R</u>eplication <u>seq</u>uencing), that combines library screening and sequencing. As a source organism for screening, we selected *S*. 6803 (library A), which contains seven plasmids in addition to the chromosome: there are at least seven replicons that can co-exist. *S*. 7942 was selected as a host cyanobacteria to be transformed with plasmid-based *S*. 6803 genomic library for three reasons: avoiding chromosome-library recombination, having extremely high transforming ability and containing only one large plasmid pANL. The *S*. 6803 genomic library (library A) was constructed on an *E. coli*-derived plasmid and introduced into *S*. 7942. As a result, we obtained more than 300 transformants. DNAs extracted from *S*. 7942 transformants were pooled as library B, which contain the autonomous replication sequence as well as *S*. 7942 genomic DNA. To isolate plasmids that replicated independently of chromosomal integration in the *S*. 7942 cells, library B was again introduced into *E. coli*, and only the plasmid DNAs were pooled as library C. At this stage, fourteen of *E. coli* colonies were picked from transformants of library B and the inserts were sequenced by Sanger sequencing. The results showed that all colonies contained the region surrounding ORF B of pCC5.2, a plasmid contained in *S*. 6803 (Supplemental Table S3).

Next, a comprehensive sequencing analysis was performed to reveal the genomic region in the libraries. The results showed that pCC5.2 occupied a large proportion in spite of its relatively short DNA length in the *S*. 6803 genomic library A (Figure 1A and Supplemental Table S2), suggesting the high copy number of pCC5.2 in *S*. 6803 cells, as reported previously (Jin et al., 2018). The fact that only a few sites were recognized by *Sau3AI* in pCC2.4 and pCB2.3 may have contributed to their relatively small proportion in the library A. Compared to the library A, the reads population of pCC5.2 significantly increased in library B and 99.8% of the reads were occupied by pCC5.2 in library C (Figure 1A and Supplementary Table S2). We mapped the sequence reads to pCC5.2 to identify the region required for replication, and observed that the region containing ORF B were concentrated in the sequence reads from libraries B and C (Figure 1B). The ORF B and the flanking region of pCC5.2 (3,242 nt) have been reported to function as a replicon to support plasmid replication in *S*. 6803 (Xu and McFadden, 1997) (Jin et al., 2018). Our study suggested that this region functions as a replicon to support plasmid replication in heterologous cyanobacterium *S*. 7942.

### 3.2 Phylogenetic analysis of ORF B in pCC5.2 (CyRepA2)

To characterize ORF B in pCC5.2, we performed a sequence-based analysis of the conservation and structure of ORF B. Analysis of the SMART and Pfam databases showed that ORF B has DUF3854 domain in its N-terminal region. The BLAST results revealed that DUF3854-containing proteins are widely conserved among cyanobacteria, not only *Synechococcales*, containing *S*. 6803 and *S*. 7942, but also *Oscillatoriophycideae, Nostocaceae, Pleurocapsales, Pseudanabaenales*, and *Chroococcidiopsidales* (Figure 2A). The DUF3854-containing proteins formed a distinctly different group from the pUH/pANS Rep and the RSF1010 RepC and were classified into two main groups. The larger group contains Slr7037 and SYNPCC7002_B0001 proteins encoded in pSYSA (*S*. 6803) and pAQ1 (*S*. 7002), which functions as a Rep in each organism (Miyasaka et al., 1998) (personal communication from Prof. Wolfgang Hess). ORF B belongs to the smaller group, containing proteins in *Microcystis, Crocosphaera*, and *Synechocystis*. Although Slr7037 and ORF B are classified into different clades, the predicted structures of these proteins are very similar, having a N-terminal DUF3854 domain and a C-terminal DEXDc domain connected by alpha-helixes (Figure 2B). DEXDc domains have been found in proteins with DNA helicase activity, and the function of the DUF3854 domain has been suggested to be related to the Toprim domain conserved in DNA primase (Honda et al., 2006), although its function remains unknown. Since the group containing Slr7037 is more common in cyanobacteria than the group containing ORF B, we designated Slr7037 as CyRepA1 (Cyanobacterial Rep protein A), and named ORF B as CyRepA2, respectively, and conducted further analysis.

### 3.3 Construction of autonomous replication plasmid pYS1 in *S*. 7942

To evaluate the replication activity of CyRepA2 in *S*. 7942, the expression vector pYS1C-GFP (Supplementary Figure S1) was constructed based on a plasmid containing the minimal region responsible for autonomous replication, obtained from the screening (1174-4683 nt in pCC5.2), and introduced into *S*. 7942 cells. pYS1C-GFP possesses *GFP* gene under the *trc* promoter and *lacI* gene, functionable in *S*. 7942, and thus can express GFP in an IPTG-dependent manner. After 48 h of pre-incubation, cells were cultivated for an additional 24 h with or without final 1 mM IPTG and used for analysis. Fluorescence microscopy exhibited an IPTG-dependent GFP fluorescence in *S*. 7942 carrying pYS1C-GFP (Figure 3A). In order to confirm the structure of pYS1C-GFP in *S*. 7942 cells, DNA was prepared from the cells harboring pYS1C-GFP and introduced into *E. coli*. Plasmids were extracted from the resulting *E. coli* transformants and compared with the initial pYS1C-GFP plasmid by restriction enzyme digestion (Supplementary Figure S2). The results proved that both plasmids were identical, suggesting that pYS1C-GFP was maintained as plasmid in *S*. 7942 cells.

To compare the utility of pYS1C-GFP with our previous expression system integrated into the chromosomal neutral site, we constructed the integration plasmid pNSG carrying *GFP* gene. The NS-GFP strain, which express GFP from chromosomal neutral site in an IPTG-dependent manner, was obtained by the transformation of *S*. 7942 with pNSG and the GFP expression levels were compared by fluorescence microscopy, flowcytometry, and western blot analysis. Microscopy showed that GFP fluorescence of the cells carrying pYS1C-GFP were significantly stronger than that of NS-GFP cells (Figure 3A), although some cells without GFP fluorescence were observed. The expression levels of GFP were quantified and compared by flowcytometry and western blotting analysis, showing that its expression in cells carrying pYS1C-GFP was about 10-fold higher than that in NS-GFP cells, and was solely dependent on IPTG (Figure 3B, C). These results indicated that the expression system using pYS1 harboring CyRepA2 is more suitable for overexpression compared to that integrated into the chromosomes in *S*. 7942.

The stability of pYS1C-GFP and the expression level of GFP in *S*. 6803 were also tested using the same approaches to *S*. 7942, because a use of pCC5.2 as an expression vector has been reported in *S*. 6803 cells (Jin et al., 2018). Consistent with the previous study, we confirmed that pYS1C-GFP is maintained autonomously as a plasmid within *S*. 6803 cell (Supplementary Figure S2). Although 5-fold of GFP expression was observed with the IPTG addition in *S*. 6803, in contrast to *S*. 7942, a leaky expression was also detected in the absence of IPTG (Supplementary Figure S3). In addition, PCR analysis indicated that the *S*. 6803 pYS1C-GFP transformants contained endogenous pCC5.2 as well, suggesting an instability of pYS1C-GFP (Supplementary Figure S4). Using more tightly regulated promoter for pYS1 in *S*. 6803 would achieve more precise expression and stable maintenance of this plasmid.

### 3.4 Compatibility of pYS and endogenous plasmid-derived vector in *S*. 7942

Phylogenetic analysis revealed that the sequence of CyRepA2 differs from that of the RepA protein encoded in the plasmid pUH24/pANS of *S*. 7942 (Figure 2A). This fact led us to assume that pYS can be maintained along with pHU24/pANS-derived vector in *S*. 7942 cells. In order to investigate the compatibility of Rep proteins, we constructed a plasmid pEX2S-mScarlet expressing orange fluorescent protein mScarlet with the spectinomycin resistance marker gene, based on the plasmid pUC303, which derived from pHU24/pANS. As a result of transformation of *S*.7942 cells carrying pYS1C-GFP with pEX2S-mScarlet, we obtained the colonies showing the resistance to both of chloramphenicol and spectinomycin (Figure 4A). Fluorescence microscopy exhibited both GFP and mScarlet fluorescence in the *S*. 7942 transformants (Figure 4B), indicating that this strain harbors pEX2S-mScarlet with pYS1C-GFP. In addition, the GFP expression levels of the cells carrying these two plasmids were comparable to those of the cells harboring only pYS1C-GFP (Supplementary Figure S5). To check the plasmid structure in *S*. 7942 cells, DNA was extracted, introduced into *E. coli* and selected by chloramphenicol or spectinomycin. Plasmid DNAs extracted from the *E. coli* transformants were digested with restriction enzymes. The results showed that the two plasmids retain the same structure as before transformation, confirming that the two plasmids were independently maintained in *S*.7942 cells (Figure 4C).

**Figure 4.**
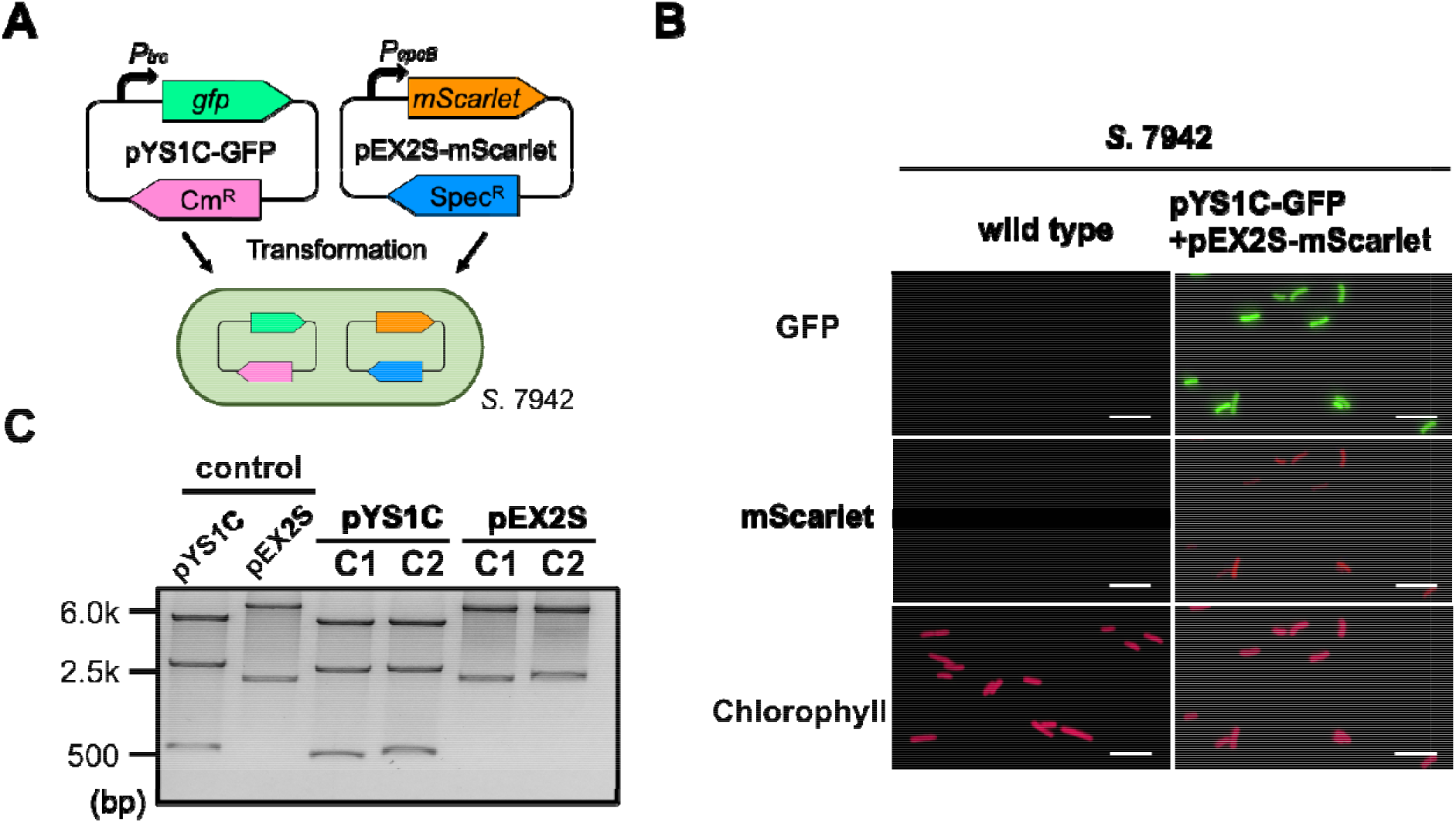
Compatibility of pYS and endogenous plasmid-derived vectors in *S*. 7942. (A) Compatibility test in *S*. 7942. pEX2S-mScarlet containing spectinomycin resistance gene (*Sp^R^*) with pHU24/pANS Rep was introduced into *S*. 7942 cell carrying pYS1C-GFP and subjected to the microscopy and DNA extraction. (B) Fluorescence microscopy. 1 mM IPTG (final conc.) was added to the culture for the induction of GFP expression in pYS1C-GFP. Images of GFP, mScarlet and chlorophyll fluorescence were shown. White bar: 10 μm. (C) Structure of the plasmids before and after the transformation to *S*. 7942. Plasmid DNAs were extracted from *S*. 7942 transformant and transformed into *E. coli*. After the extraction from *E. coli*, plasmids were digested with restriction enzymes, and compared with the plasmid before transformation to *S*. 7942 (control). *NdeI* and *PstI* (for pYS1C-GFP) and *SalI* and *NheI* (for pEX2S-mScarlet) were used for the digestion. The plasmids before transformation was used as a control. The results of two independent clones were shown as C1 and C2.

### 3.5 Development of expression vectors functioning in *S*. 7002

DUF3854-containing protein was found in pAQ1 plasmid of the model marine cyanobacterium *S*. 7002 (SYNPCC7002_B0001), which is closely related to CyRepA1 rather than CyRepA2 (Figure 2). Thus, we tested the availability of pYS1 in *S*. 7002 according to the similar procedure in both cases of *S*. 7942 and *S*. 6803. As in the previous two cyanobacteria, significant GFP expression and plasmid maintenance was observed in *S*. 7002 (Figure 5A-D and Supplementary Figure S2). To reveal the compatibility between pYS plasmid and endogenous pAQ1 in *S*. 7002 cells, we performed PCR analysis using specific primer sets for the amplification of pYS1C-GFP and pAQ1, respectively. The results showed that the pAQ1 is stably retained in the pYS1C-GFP transformant (Supplementary Figure S4), indicating that the replication system of CyRepA2-type can be co-exist with that of CyRepA1-type despite the similarity of their amino-acid sequences.

**Figure 5.**
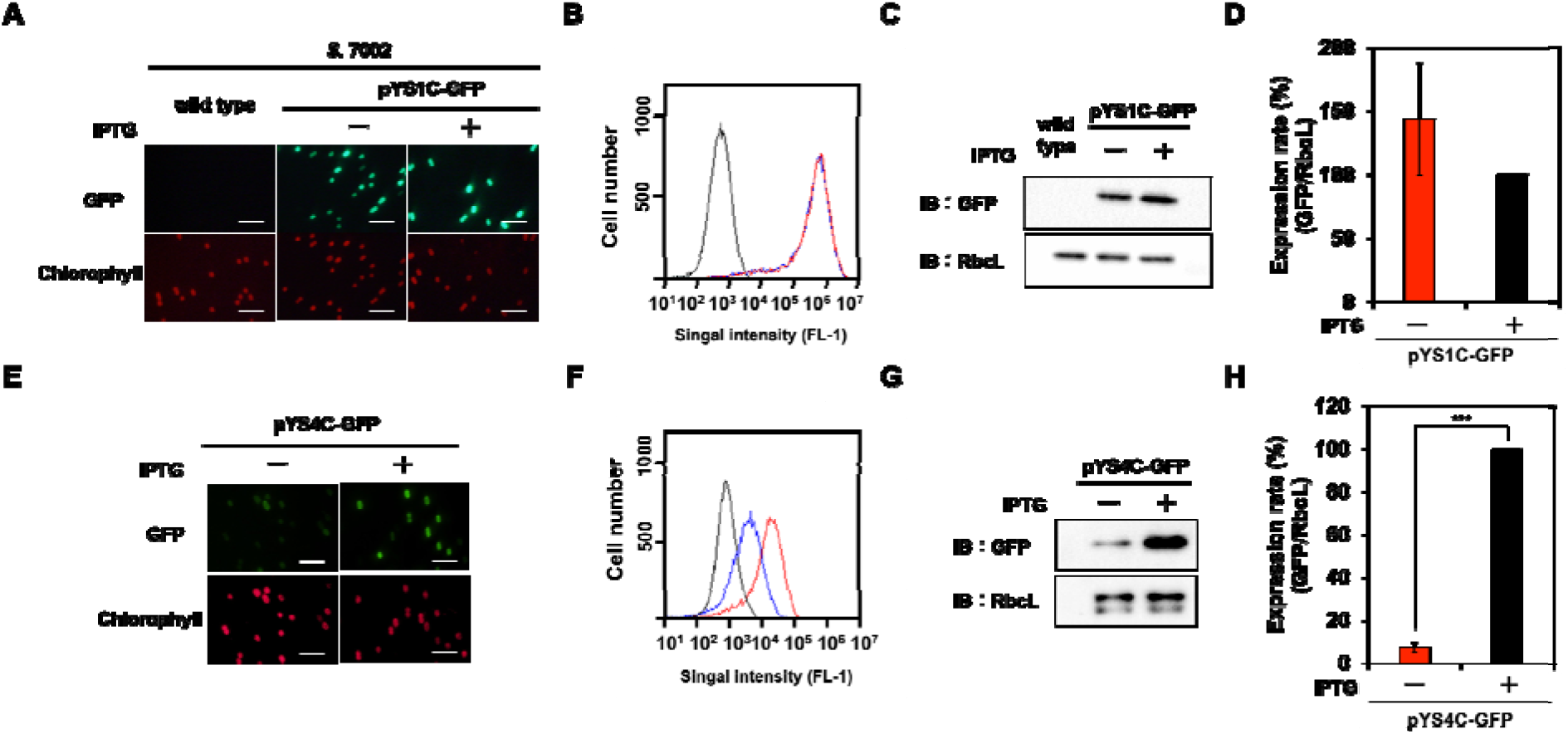
Utilization of expression plasmid pYS1 and pYS4 in *S*. 7002. The GFP expression levels in pYS1C-GFP (A-D) and pYS4C-GFP transformants (E-H) were analyzed with (+) and without (-) final 1 mM IPTG. (A, E) Fluorescence microscopy. Images of GFP and chlorophyll were shown. White bar: 10 μm (B, F) FACS analysis of GFP fluorescence. Signal intensity of FL-1 indicating GFP fluorescence in *S*. 7002 wild type (black), pYS1- or pYS4-GFP in the presence (red) and absence (blue) of IPTG were shown. (C, G) Western blotting analysis. The protein extracts obtained from *S*. 7002 cells were subjected to SDS-PAGE and analyzed by western blotting using antibodies against GFP and control RbcL used as an internal control. (D, H) Comparison of the GFP signal. The signals intensity of GFP obtained from western blotting analysis were normalized with those of RbcL and the ratio of GFP signal at presence of IPTG was set to 100. Bars represent mean ± SEM (*n* = 3) (***p < 0.005).

In contrast to the cases of *S*. 7942 and *S*. 6803, there was no difference between the expressions (both fluorescence and amount) of GFP proteins in the presence and absence of IPTG in *S*. 7002 (Figure 5A-D), indicating that the repression of *trc* promoter by LacI does not work at all in *S*. 7002 as reported in previously (Markley et al., 2015). We noted a broad distribution of the weaker fluorescence signal in the FACS profile of *S*. 7002 (Figure 5B), suggesting that there are differences in the GFP expression level varies in each cell, which is consistent with the GFP fluorescence observed in the microscopy.

To improve the induction system of pYS1C-GFP in *S*. 7002 cells, the *trc* promoter was replaced with *clac143* promoter, which has been shown to enable an IPTG-dependent regulation in *S*. 7002 (Markley et al., 2015). The resulting plasmid, named pYS4C-GFP, was introduced into *S*. 7002, and the GFP expression was tested in the presence or absence of IPTG. SDS-western blotting analysis demonstrated that the expression level of GFP can be controlled by IPTG (Figure 5E-H). Comparison of GFP fluorescence intensity of pYS1C-GFP and pYS4C-GFP strains in the presence of IPTG by flowcytometry showed that the fluorescence of the cells carrying pYS1C-GFP is more than one order of magnitude higher than in those carrying pYS4C-GFP, indicating that pYS1 can be used as an over-expression system and pYS4, on the other hand, as a strict control system (Figure 5B, F).

### 3.6 Utilization of pYS vector in *A*. 7120

To further test the utility of the pYS plasmid, we introduced it via conjugation into a filamentous nitrogen-fixing cyanobacteria *A*. 7120 which belongs to *Nostocaceae*, a phylogenetic group distinct from the *Synechococcales* including *S*. 6803, *S*. 7942 and *S*. 7002. For the conjugation transport of pYS plasmid to *A*. 7120, which requires a helper plasmid pRL carrying *cat* marker gene, we replaced the *cat* gene in pYS1C- and pYS4C-GFP with spectinomycin resistant marker gene and named the resulting plasmids as pYS1S-GFP and pYS4S-GFP, respectively. We successfully isolated *A*. 7120 transconjugants carrying pYS4S-GFP, suggesting that this plasmid can be transported to *A*. 7120 cells by the conjugation, although the transconjugant carrying pYS1S-GFP was not obtained. This may be due to an over-expression of GFP by the *trc* promoter, as in *S*. 7002, which could be too strong and cytotoxic in the *A*. 7120 cells. In the pYS4S-GFP transconjugants, GFP fluorescence was observed in every filamentous cell and the structure of pYS4S-GFP was identical to that of the plasmid before transconjugation (Figure 6A and Supplementary Figure S2), suggesting the pYS4-derived vector was maintained stably in *A*. 7120 cells, although, in contrast to *S*. 7002, there was no difference between the GFP expression in the presence and absence of IPTG (Figure 6B, C).

**Figure 6.**
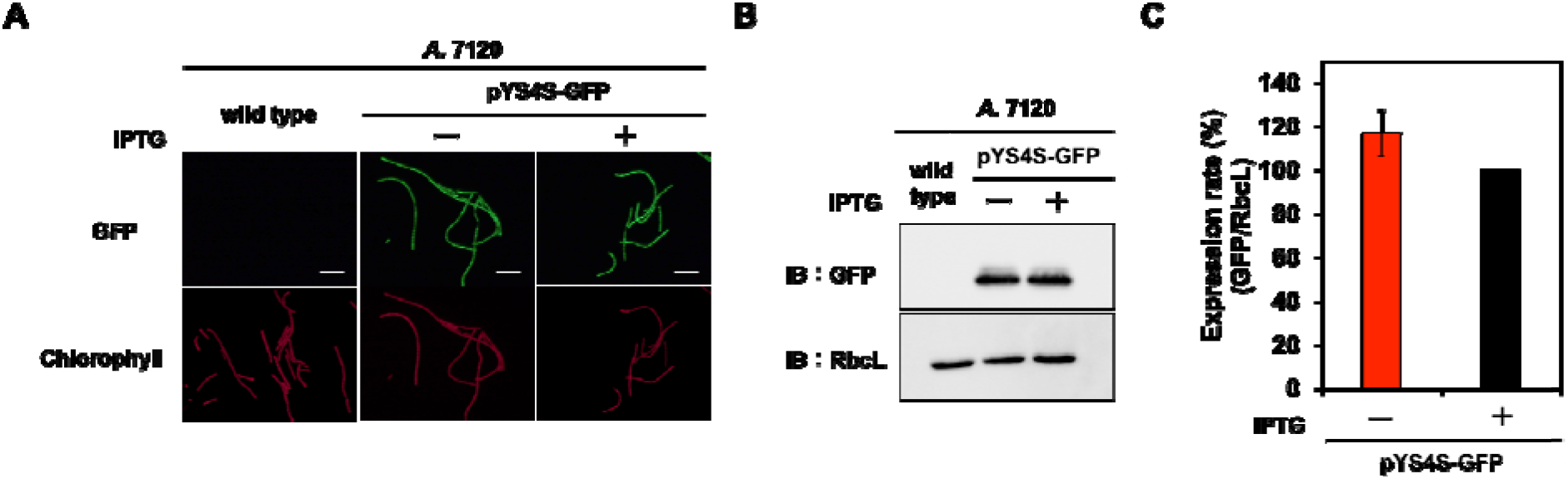
Utilization of pYS in *A*. 7120. The GFP expression levels in pYS4S-GFP transconjugants were analyzed with (+) and without (-) final 1 mM IPTG. (A) Fluorescence microscopy. Images of GFP and chlorophyll were shown. White bar: 20 μm (B) Western blotting analysis. The protein extracts obtained from *A*. 7120 cells were subjected to SDS-PAGE and analyzed by western blotting using antibodies against GFP and control RbcL used as an internal control. (C) The comparison of the GFP signal. The signals of GFP obtained from western blotting analysis were normalized with that of RbcL signal and the ratio of GFP signal at presence of IPTG was set to 100. Bars represent mean ± SEM (*n* = 3).

## 4 Discussion

Cyanobacteria, which grow with CO2 absorption through photosynthesis, are promising hosts for carbon-neutral material production. However, even with model cyanobacteria such as *S*. 7942 and *S*. 6803, only a limited number of vectors are available for genetic engineering. We have newly established AR-seq as a novel screening method for autonomous replication regions and found that the CyRepA2-containing region derived from *S*. 6803 can be used for overexpression vectors in a wide range of cyanobacterial species.

In several organisms, not only bacteria but also eukaryotic microorganisms, autonomous replication origins have been explored using genome libraries to identify replication initiation sites and factors involved in DNA replication (Yasuda and Hirota, 1977) (Hsiao and Carbon, 1979) (Rochaix et al., 1984). AR-seq, combination methodology of the traditional approach with NGS technology, is a powerful tool for finding autonomous replication regions in the genome, including regions that function as DNA replication start sites as well as genes encoding replication factors. Our screening of the *S*. 6803 genomic library revealed that the ORF of CyRepA2 and its upstream region are required for autonomous replication (Figure 1B and Supplementary Table S3), which is consistent with the previous study (Jin et al., 2018). It would be possible to identify new Rep and autonomous replication regions by changing the genomic library and host organisms used for AR-seq. Given its versatility, AR-seq could be used to identify autonomous replication initiation regions not only in cyanobacteria, but also in a variety of other organisms.

It has been reported that pCC5.2 exhibits rolling-circle replication (RCR) (Xu and McFadden, 1997). RCR represents one of the simplest initiation strategies, starting from a nick on one parental strand generated around inverted repeat sequences, which functions as a primer for DNA replication initiation of RCR plasmids. The inverted repeats are found in the ORF of CyRepA2 and have been proposed to be a replication initiation site in pCC5.2 (Xu and McFadden, 1997). Many of the RCR plasmids have been shown to replicate in species, genera, or even phyla other than those from which they were isolated. The simplicity of the RCR initiation, with only the plasmid-encoded Rep protein participating in origin recognition and priming of the leading strand synthesis, is thought to underlie the usual promiscuity of RCR plasmids (del Solar et al., 2014). Consistent with these observations, the pYS plasmid constructed in this study based on pCC5.2 also exhibited a wide host range, similar to the RCR-employing pSOMA plasmids derived from pCA2.4 and pCB2.4 (Opel et al., 2022).

RCR plasmids appear to be suitable for expression due to their generally small size, high copy number, and few specific factors involved in replication. RCR plasmids are commonly used in the industrial microorganism *Corynebacterium glutamicum* (Pátek and Nešvera, 2013). For example, RCR vectors have been used to overexpress the *pyc* gene to yield large amounts of glutamate, lysine and threonine (Peters-Wendisch et al., 2001). Our results showed that pYS can be used for gene overexpression in several cyanobacteria. Especially in *S*. 7942, GFP expression was 10-fold higher than in chromosome-based expression systems, which can be tightly controlled by IPTG. These results strongly suggest that the pYS vectors constructed in this study are exceptional genetic-engineering tools for potential uses at the industrial level in cyanobacteria.

Other utilities of the RCR plasmids include their usage in combination with other plasmids. In this study we showed that pYS derived from pCC5.2 can be harbored simultaneously with the pEX2, derived from an endogenous plasmid in *S*. 7942. Furthermore, we also demonstrated that the pAQ1 can be maintained along with pYS1C-GFP in *S*. 7002 cells (Supplementary Figure S4), indicating that the pYS can be used together with pAQ-based expression systems. It has been known that the RCR plasmids can be maintained together with other plasmids in *S*. 6803 cells: pCC5.2-derived vector pSCB can coexist with the RSF1010-based vector in the same host cell (Jin et al., 2018), and pSOMA10 derived from pCA2.4 can be harbored with pSOMA16 (derived pCB2.4) or RSF1010-based vectors (Opel et al., 2022). The utilization of plasmids in various combinations would allow complex genetic modifications in cyanobacteria, such as metabolic engineering. Further basic research on these RCR plasmids will enable the development of more suitable plasmids for material production in cyanobacteria.

The structure prediction indicates that CyRepA is a multi-domain protein (Figure 2B). The fact that CyRepA2 has a DUF3854 domain related to the Toprim domain observed in DNA primase, even though RCR does not require primer RNA synthesis, is interesting, considering the evolution of CyRepA protein. The predicted structure of CyRepA2 is very similar to that of CyRepA1 (Figure 2B). However, the two CyRepAs are compatible in *S*. 6803 cells, because pSYSA (coding CyRepA1) and pCC5.2 or pYS1C-GFP (coding CyRepA2) coexist in *S*. 6803 cells (Supplementary Figure S4). In *S*. 7002, the pYS1C-GFP transformant stably retains the pAQ1 plasmid carrying CyRepA1 homolog SYNPCC7002_B0001 (Supplementary Figure S4). These observations indicate that CyRepA1 and CyRepA2 can function independently as replication factors in cyanobacteria cells. While the size of pSYSA is about 20 times larger than pCC5.2, the copy number of pSYSA in *S*. 6803 cells is clearly lower than that of pCC5.2, suggesting that CyRepA1 and CyRepA2 have different properties as replication factors (Nagy et al., 2021). The predicted structures provide important insights into the functional differences between CyRepA1 and CyRepA2. Especially in the N-terminal region, a clear structural difference between the two is observed (Figure 2B). Since CyRepA2 is found in some closely related cyanobacteria such as *Synechocystis*, *Microcystis*, and *Crocosphaera* (Figure 2A), it may have originated from CyRepA1 in the common ancestor of these species. Future comparisons of CyRepA1 and CyRepA2 will reveal functional differences, such as host range and replication activity in cyanobacteria.

For additional requirements for specific article types and further information please refer to <u>Author Guidelines</u>.

## 6 Acknowledgments

We are grateful to Prof. Wolfgang Hess for providing valuable observations on the function of the Slr7037 protein. We thank Prof. Daisuke Umeno and Dr. Ryudo Ohbayashi for providing the plasmids for construct pYS1C-GFP and pEX2S-mScarlet, respectively. We sincerely thank Dr. Shigeki Ehira for advising us on the transformation of *A*.7120. We also appreciate to Yuka Kinouchi for her technical assistance.

