## Supplementary Data for "Exploration of the autonomous replication region and its utilization for expression vectors in cyanobacteria"

**Supplemental Table S1.**

Oligonucleotide primers used in this study.

Primer name	Sequence (5′ to 3′)*^a^*

*The construction of S. 6803 genomic library*

F1	CAACATACGAGCCGGAAGCATAAAGT

R2	GCGGATCCCTGTCAGACCAAGTTTACTCATATATACTTTAGATTGA

F3	GCGGATCCAGAATAAATAAATCCTGGTGTCCCTGTTGA

R4	CCGGCTCGTATGTTGTTACGCCCCGCCCTGCCA

*Plasmid and strain construction*

F5	AGAATAAATAAATCCTGGTGTCCCTGTTG

R6	GATCGTCGTCGCTCAAAAAGAGACTAATAAC

F7	TGAGCGACGACGATCCCAGGCATCAAATAAAACGA

R8	AGCGTCGAGATCCCGGACACCATCGAATGGCGC

F9	CGGGATCTCGACGCTCTCCCTTATGCGACT

R10	GATAACCTCCTAAATTGTTATCCGCTCACAATT

F11	ATTTAGGAGGTTATCATGGAATTCAGTAAAGGAGAAGAACTTTTC

R12	GTATTACTAGTAAGCTTATTTGTATAGTTCATCCATGCCATGTG

F13	GCTTACTAGTAATACTGCAGAGAGA

R14	GGATTTATTTATTCTCACATTTCCCCGAAAAGTG

F15 TGTAATTGACATAAGTCCCATCACCGTTGTATAAATGTGTGGAATTGTGAGCGGATAACAATTTCACACAATGGAATTCAGTAAAGGAGAAGAACTTTTCACTG

R16	TGGGACTTATGTCAATTACATCTTGTTAATTTTATTCCTGCTTTTTTGTTAAGAATTCCGAATTGTGAGCGCTCACAATTCGGGCTCATGAGCGCTTGTTTC

F17	CAACATACGAGCCGGAAGCATAAAGTGTAA

R18	CACATTTCCCCGAAAAGTGCCACCTGA

F19	TTTCGGGGAAATGTGCGCAGCGGTGGTAACGGCG

R20	TGCTTCCGGCTCGTATGTTGTTATTTGCCGACTACCTTGG

F21	GGCGTCGACAGTAAAGGAGAAGAACTTT

R22	GGCAAGCTTTTATTTGTATAGTTCATCC

F23	AACACCTTCGGGAGAGCCTGTTAACACT

R24	CATAGTCGAGTTACGGATCTGCAAGTCAACAGCCGCG

F25	AACTCTTAGATCTGCCACCGCCGGACATCAGCGCTAG

R26	TCTCCCGAAGGTGTTTCAAACATGAGAATTACAACTTATATCG

F27	CGTAACTCGACTATGCTTGTAAACCGT

R28	GCAGATCTAAGAGTTTGTAGAAACGCAAAAAGGCC

*Plasmid PCR*

F29	CCGGCTCGTATAATGTGTGG

R30	GAGCAACTGACTGAAATGCCTCA

F31	ATTGTCTGTAGGTAAGTTTTTTAGCGTC

R32	TGAGAAGACTATCCTGCCCAAC

F33	TTGCTCCCGCCACATGGT

R34	GCCATTCTTTTCCTCCATCACTGCGGTGG

F35	TGATCGAAATACTCGTTGTGCAG

R36	GAAACAGAAAATCTAAAGACCAACCCG

Supplementary Table S2.

|  | Library A | Library B | Library C |
| --- | --- | --- | --- |
| chromosome (3,570 kbp) | 80531 (61.851%) | 3734 (5.853%) | 1579 (0.161%) |
| pSYSM (120 kbp) | 4722 (3.627%) | 52 (0.082%) | 18 (0.002%) |
| pSYSX (106 kbp) | 11471 (8.810%) | 119 (0.187%) | 128 (0.013%) |
| pSYSA (103 kbp) | 4666 (3.584%) | 24 (0.038%) | 12 (0.001%) |
| pSYSG (44.3 kbp) | 960 (0.737%) | 20 (0.031%) | 14 (0.001%) |
| pCA2.4 (2.4 kbp) | 1265 (0.972%) | 12 (0.019%) | 16 (0.002%) |
| pCB2.4 (2.4 kbp) | 517 (0.397%) | 9 (0.014%) | 9 (0.001%) |
| **pCC5.2 (5.2 kbp)** | **26069 (20.022%)** | **59821 (93.777%)** | **981084 (99.819%)** |
| Total | 130201 (100%) | 63791 (100%) | 982860 (100%) |

Sequencing results of *S*. 6803 genomic libraries. Sequencing reads in each library were mapped to the *S*. 6803 genome (chromosome and 7 plasmids, pSYSM, pSYSX, pSYSA, pSYSG, pCA2.4, pCB2.4, and pCC5.2). The number of reads and ration in each library were shown.

Supplementary Table S3.

| Clone number | Name | Genbank ID | Strat point | End point | Length (bp) |
| --- | --- | --- | --- | --- | --- |
| 1 | pCC5.2 | CP003272.1 | 1174 | 4726 | 3553 |
| 2 | pCC5.2 | CP003272.1 | 1174 | 5133 | 3960 |
| 3* | pCC5.2 | CP003272.1 | 1174 | 4683 | 3510 |
| 4 | pCC5.2 | CP003272.1 | 4763 | 1174 | 3590 |
| 5 | pCC5.2 | CP003272.1 | 1174 | 4763 | 3590 |
| 6 | pCC5.2 | CP003272.1 | 895 | 5110 | 4216 |
| 7 | pCC5.2 | CP003272.1 | 895 | 4674 | 3780 |
| 8 | pCC5.2 | CP003272.1 | 1174 | 5159 | 3986 |
| 9 | pCC5.2 | CP003272.1 | 895 | 5087 | 4193 |
| 10 | pCC5.2 | CP003272.1 | 1174 | 5159 | 3986 |
| 11 | pCC5.2 | CP003272.1 | 5159 | 1174 | 3986 |
| 12 | pCC5.2 | CP003272.1 | 1174 | 4763 | 3590 |
| 13 | pCC5.2 | CP003272.1 | 895 | 5159 | 4265 |
| 14 | pCC5.2 | CP003272.1 | 1177 | 1202 | 5188 |

Fourteen of *E. coli* colonies were selected from transformants of Library B and the sequences of their insert regions were determined by Sanger sequencing. All sequenced regions were matched to the plasmid pCC5.2 in *S*. 6803. The base numbers in the pCC5.2 sequence (Genbank ID: CP003272.1) at the start point and end point of the insert regions were shown along with the length of the inserts.

*Clone 3 contains minimum insert region was used for constructing expression vector pYS.

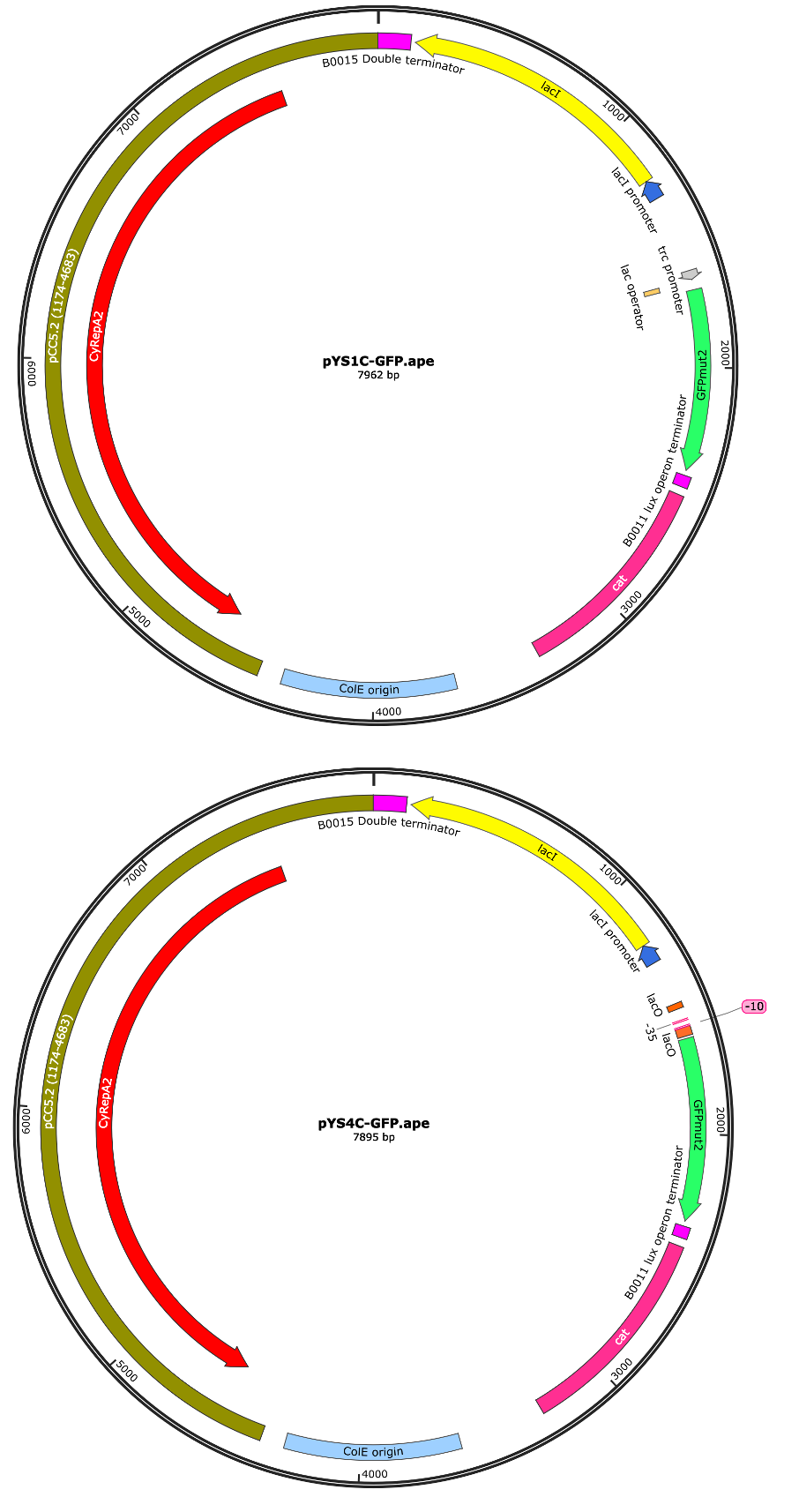

**Supplementary Figure S1. The vector map of pYS1C-GFP and pYS4C-GFP.**

The expression vectors pYS1C-GFP and pYS4C-GFP were constructed from plasmids containing the minimum region of autonomous replication activity (1174-4683 nt including *CyRepA2* gene of pCC5.2), *cat* gene and *ColE* region obtained by screening. To monitor the expression level in an IPTG dependent manner, GFP gene was placed under the *trc* promoter (pYS1) or the *cLac143* promoter (pYS4), containing *lacO* operator, together with the repressor *lacI* gene. The image of vector map was drawn using SnapGene software. The vector sequence can be obtained from Supplementary Materials.

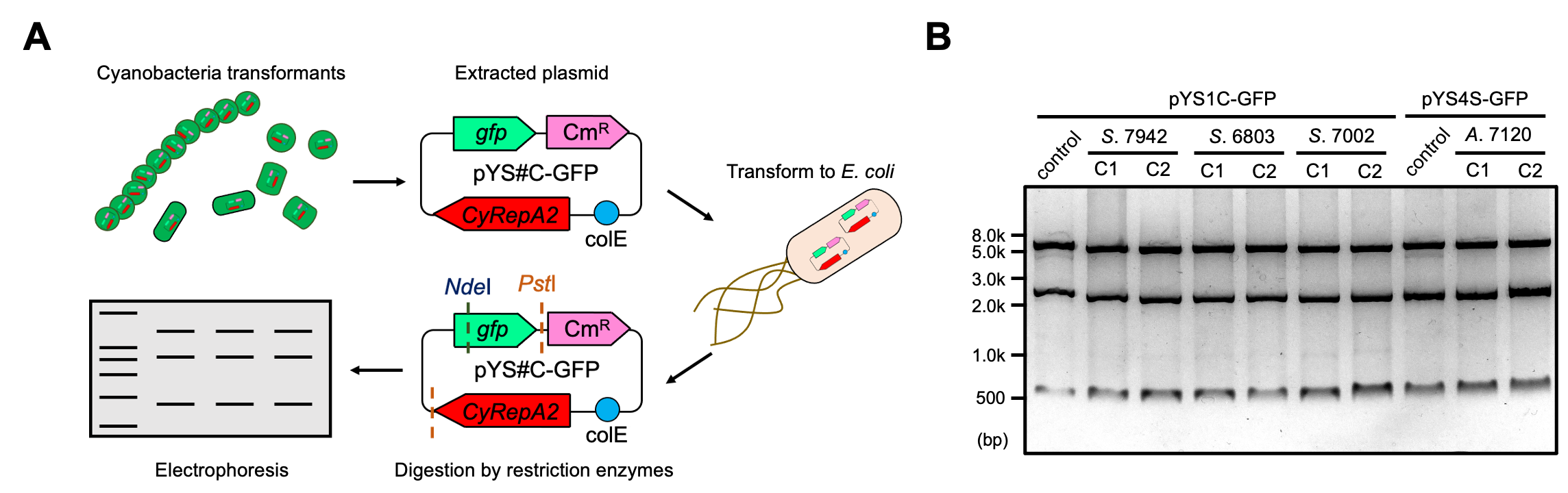

**Supplementary Figure S2. Plasmid structure of pYS in cyanobacteria cells.**

(A) Scheme of the analysis of plasmid structure. To determine whether plasmids are maintained in cyanobacterial cells in a circular structure, DNA extracted from cyanobacteria transformants (*S*. 7942, *S*. 6803, *S*. 7002, and *A*. 7120), carrying pYS1C-GFP or pYS4S-GFP, were introduced into *E. coli*. After the plasmid extraction from the *E. coli* cells, plasmids were digested with restriction enzymes, and compared to the plasmids before transformation to cyanobacteria. (B) Electrophoresis image of plasmids digested by the restriction enzymes. The plasmids before transformation was used as a control. The results of two independent clones were shown as C1 and C2.

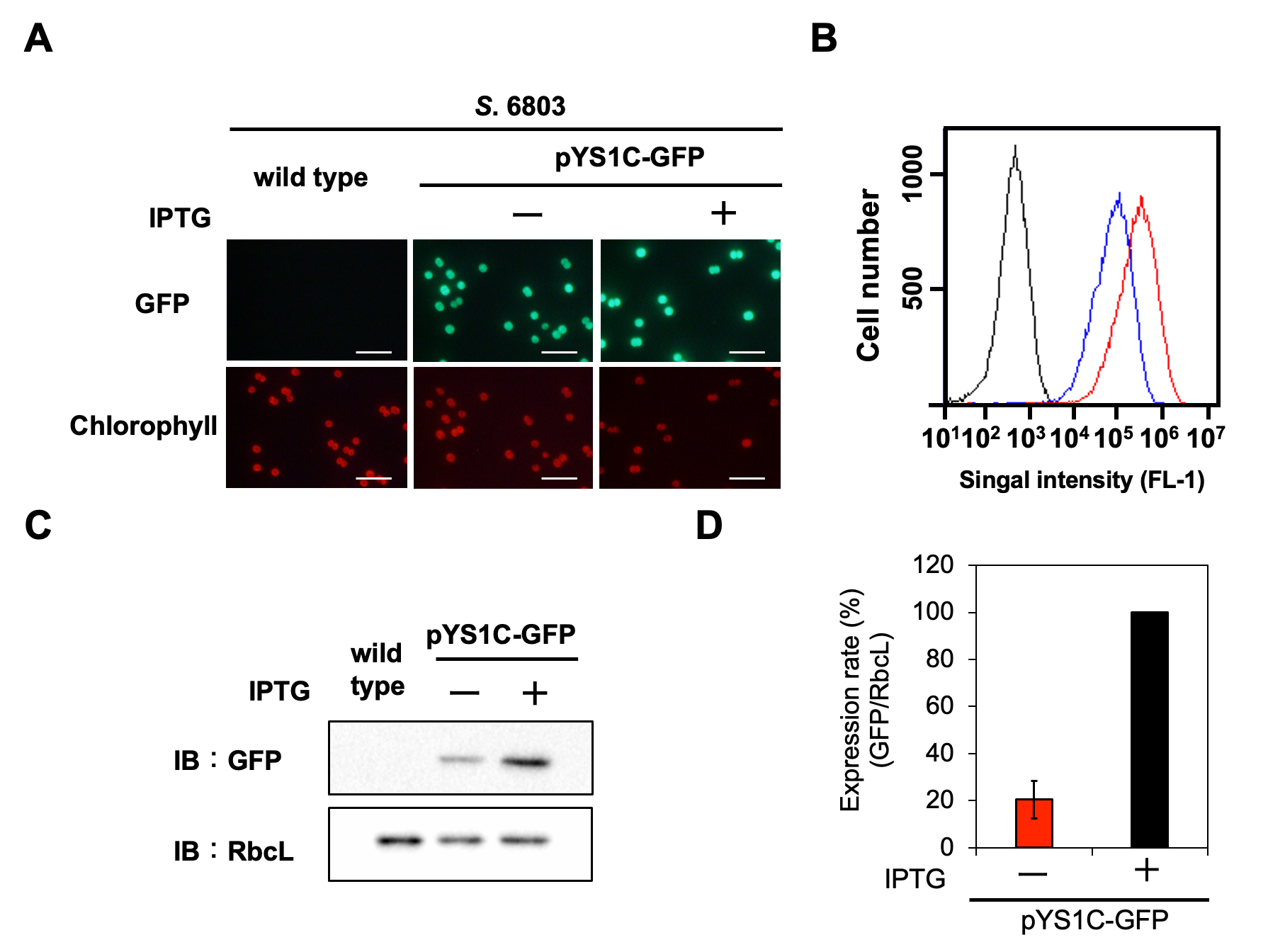

**Supplementary Figure S3. Utilization of pYS1 in *S*. 6803.**

The GFP expression levels in pYS1C-GFP were analyzed with (+) and without (-) final 1 mM IPTG. (A) Fluorescence microscopy. Images of GFP and chlorophyll were shown. White bar: 10 μm (B) FACS analysis of GFP fluorescence. Signal intensity of FL1 indicating GFP fluorescence in *S*. 6803 wild type (black) and pYS1C-GFP transformants in the presence (red) and absence (blue) of IPTG were shown. (C) Western blotting analysis. The protein extracts obtained from *S*. 6803 cells were subjected to SDS-PAGE and analyzed by western blotting using antibodies against GFP and control RbcL used as an internal control. (D) Comparison of the GFP signal. The signals intensity of GFP obtained from western blotting analysis were normalized with those of RbcL and the ratio of GFP signal at presence of IPTG was set to 100. Bars represent mean ± SEM (*n* = 3).

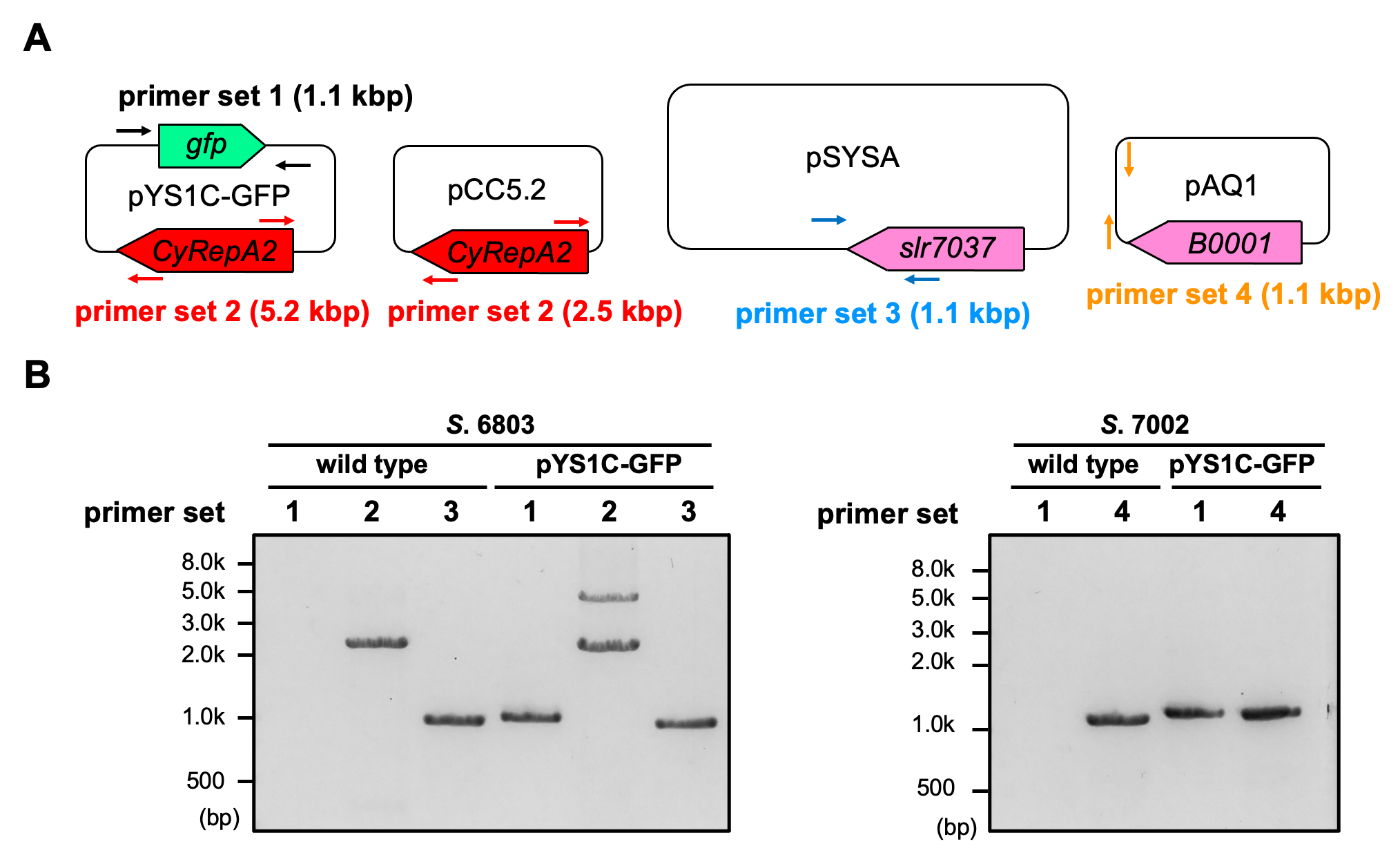

**Supplementary Figure S4.　Compatibility of pYS and endogenous plasmids in *S*. 6803 and *S*.7002.**

(A) A schematic diagram of the analyzed plasmids and the primers used for the PCR analysis. (B) electrophoresis image of PCR products. DNA containing plasmids were extracted from wild type and pYS1C-GFP transformants in *S*. 6803 and *S*.7002 and were PCR-amplified with the appropriate primer sets. pYS1C-GFP: primersF29 and R 30, pCC5.2: primers F 31 and R32, pSYSA: primers F33 and R34, pAQ1: primers (F35 and R36 (Supplementary Table S1).

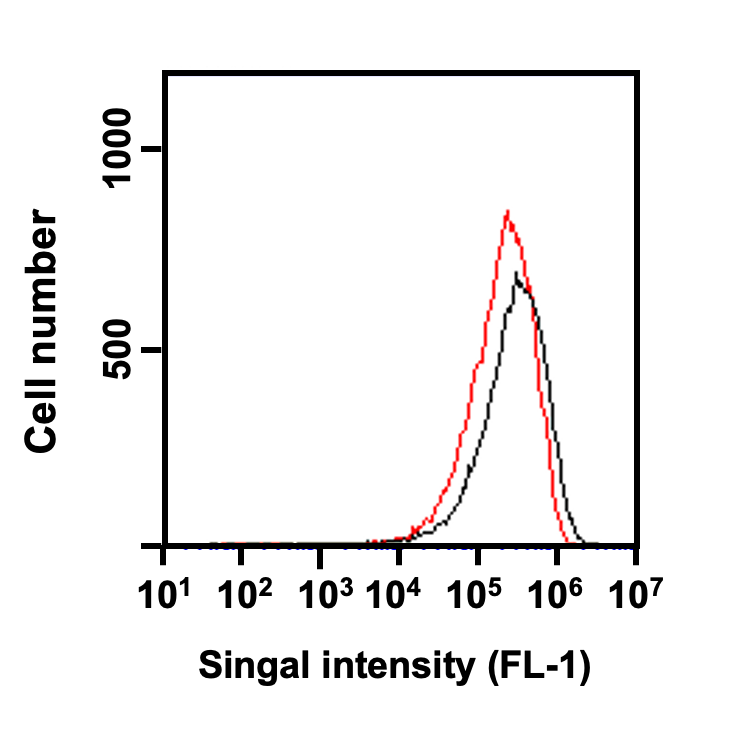

**Supplementary Figure S5. Effect of maintenance of the secondary plasmid on GFP expression lovels in pYS plasmid.**

GFP fluorescence (FL1) were compared between. *S*. 7942 cells harboring pYS1C-GFP (black) and pEX2S-mScarlet in attrition to pYS1C-GFP (red). Final 1 mM IPTG was added to induce GFP expression.

**Supplementary material Data 1**

CCAGGCATCAAATAAAACGAAAGGCTCAGTCGAAAGACTGGGCCTTTCGTTTTATCTGTTGTTTGTCGGTGAACGCTCTCTACTAGAGTCACACTGGCTCACCTTCGGGTGGGCCTTTCTGCGTTTATACATTAATTGCGTTGCGCTCACTGCCCGCTTTCCAGTCGGGAAACCTGTCGTGCCAGCTGCATTAATGAATCGGCCAACGCGCGGGGAGAGGCGGTTTGCGTATTGGGCGCCAGGGTGGTTTTTCTTTTCACCAGTGAGACGGGCAACAGCTGATTGCCCTTCACCGCCTGGCCCTGAGAGAGTTGCAGCAAGCGGTCCACGCTGGTTTGCCCCAGCAGGCGAAAATCCTGTTTGATGGTGGTTAACGGCGGGATATAACATGAGCTGTCTTCGGTATCGTCGTATCCCACTACCGAGATATCCGCACCAACGCGCAGCCCGGACTCGGTAATGGCGCGCATTGCGCCCAGCGCCATCTGATCGTTGGCAACCAGCATCGCAGTGGGAACGATGCCCTCATTCAGCATTTGCATGGTTTGTTGAAAACCGGACATGGCACTCCAGTCGCCTTCCCGTTCCGCTATCGGCTGAATTTGATTGCGAGTGAGATATTTATGCCAGCCAGCCAGACGCAGACGCGCCGAGACAGAACTTAATGGGCCCGCTAACAGCGCGATTTGCTGGTGACCCAATGCGACCAGATGCTCCACGCCCAGTCGCGTACCGTCTTCATGGGAGAAAATAATACTGTTGATGGGTGTCTGGTCAGAGACATCAAGAAATAACGCCGGAACATTAGTGCAGGCAGCTTCCACAGCAATGGCATCCTGGTCATCCAGCGGATAGTTAATGATCAGCCCACTGACGCGTTGCGCGAGAAGATTGTGCACCGCCGCTTTACAGGCTTCGACGCCGCTTCGTTCTACCATCGACACCACCACGCTGGCACCCAGTTGATCGGCGCGAGATTTAATCGCCGCGACAATTTGCGACGGCGCGTGCAGGGCCAGACTGGAGGTGGCAACGCCAATCAGCAACGACTGTTTGCCCGCCAGTTGTTGTGCCACGCGGTTGGGAATGTAATTCAGCTCCGCCATCGCCGCTTCCACTTTTTCCCGCGTTTTCGCAGAAACGTGGCTGGCCTGGTTCACCACGCGGGAAACGGTCTGATAAGAGACACCGGCATACTCTGCGACATCGTATAACGTTACTGGTTTCACATTCACCACCCTGAATTGACTCTCTTCCGGGCGCTATCATGCCATACCGCGAAAGGTTTTGCGCCATTCGATGGTGTCCGGGATCTCGACGCTCTCCCTTATGCGACTCCTGCATTAGGAAGCAGCCCAGTAGTAGGTTGAGGCCGTTGAGCACCGCCGCCGCAAGGAATGGTGCATGCAAGGAGATGGCGCCCAACAGTCCCCCGGCCACGGGGCCTGCCACCATACCCACGCCGAAACAAGCGCTCATGAGCCCGAAGTGGCGAGCCCGATCTTCCCCATCGGTGATGTCGGCGATATAGGCGCCAGCAACCGCACCTGTGGCGCCGGTGATGCCGGCCACGATGCGTCCGGCGTAGAGGATCGAGATCTCGGGTACCGAGCTGTTGACAATTAATCATCCGGCTCGTATAATGTGTGGAATTGTGAGCGGATAACAATTTAGGAGGTTATCATGGAATTCAGTAAAGGAGAAGAACTTTTCACTGGAGTTGTCCCAATTCTTGTTGAATTAGATGGTGATGTTAATGGGCACAAATTTTCTGTCAGTGGAGAGGGTGAAGGTGATGCAACATACGGAAAACTTACCCTTAAATTTATTTGCACTACTGGAAAACTACCTGTTCCATGGCCAACACTTGTCACTACTTTCGCGTATGGTCTTCAATGCTTTGCGAGATACCCAGATCATATGAAACAGCATGACTTTTTCAAGAGTGCCATGCCCGAAGGTTATGTACAGGAAAGAACTATATTTTTCAAAGATGACGGGAACTACAAGACACGTGCTGAAGTCAAGTTTGAAGGTGATACCCTTGTTAATAGAATCGAGTTAAAAGGTATTGATTTTAAAGAAGATGGAAACATTCTTGGACACAAATTGGAATACAACTATAACTCACACAATGTATACATCATGGCAGACAAACAAAAGAATGGAATCAAAGTTAACTTCAAAATTAGACACAACATTGAAGATGGAAGCGTTCAACTAGCAGACCATTATCAACAAAATACTCCAATTGGCGATGGCCCTGTCCTTTTACCAGACAACCATTACCTGTCCACACAATCTGCCCTTTCGAAAGATCCCAACGAAAAGAGAGACCACATGGTCCTTCTTGAGTTTGTAACAGCTGCTGGGATTACACATGGCATGGATGAACTATACAAATAAGCTTACTAGTAATACTGCAGAGAGAATATAAAAAGCCAGATTATTAATCCGGCTTTTTTATTATTTAGACGTCAGGTGGCACTTTTCGGGGAAATGTGAGAATAAATAAATCCTGGTGTCCCTGTTGATACCGGGAAGCCCTGGGCCAACTTTTGGCGAAAATGAGACGTTGATCGGCACGTAAGAGGTTCCAACTTTCACCATAATGAAATAAGATCACTACCGGGCGTATTTTTTGAGTTATCGAGATTTTCAGGAGCTAAGGAAGCTAAAATGGAGAAAAAAATCACTGGATATACCACCGTTGATATATCCCAATGGCATCGTAAAGAACATTTTGAGGCATTTCAGTCAGTTGCTCAATGTACCTATAACCAGACCGTTCAGCTGGATATTACGGCCTTTTTAAAGACCGTAAAGAAAAATAAGCACAAGTTTTATCCGGCCTTTATTCACATTCTTGCCCGCCTGATGAATGCTCATCCGGAATTCCGTATGGCAATGAAAGACGGTGAGCTGGTGATATGGGATAGTGTTCACCCTTGTTACACCGTTTTCCATGAGCAAACTGAAACGTTTTCATCGCTCTGGAGTGAATACCACGACGATTTCCGGCAGTTTCTACACATATATTCGCAAGATGTGGCGTGTTACGGTGAAAACCTGGCCTATTTCCCTAAAGGGTTTATTGAGAATATGTTTTTCGTCTCAGCCAATCCCTGGGTGAGTTTCACCAGTTTTGATTTAAACGTGGCCAATATGGACAACTTCTTCGCCCCCGTTTTCACCATGGGCAAATATTATACGCAAGGCGACAAGGTGCTGATGCCGCTGGCGATTCAGGTTCATCATGCCGTTTGTGATGGCTTCCATGTCGGCAGAATGCTTAATGAATTACAACAGTACTGCGATGAGTGGCAGGGCGGGGCGTAACAACATACGAGCCGGAAGCATAAAGTGTAAAGCCTGGGGTGCCTAATGAGTGAGCTAACTCACATTAATTGCGTTGCGCTCACTGCCCGCTTTCCAGTCGGGAAACCTGTCGTGCCAGCTGCATTAATGAATCGGCCAACGCGCGGGGAGAGGCGGTTTGCGTATTGGGCGCTCTTCCGCTTCCTCGCTCACTGACTCGCTGCGCTCGGTCGTTCGGCTGCGGCGAGCGGTATCAGCTCACTCAAAGGCGGTAATACGGTTATCCACAGAATCAGGGGATAACGCAGGAAAGAACATGTGAGCAAAAGGCCAGCAAAAGGCCAGGAACCGTAAAAAGGCCGCGTTGCTGGCGTTTTTCCATAGGCTCCGCCCCCCTGACGAGCATCACAAAAATCGACGCTCAAGTCAGAGGTGGCGAAACCCGACAGGACTATAAAGATACCAGGCGTTTCCCCCTGGAAGCTCCCTCGTGCGCTCTCCTGTTCCGACCCTGCCGCTTACCGGATACCTGTCCGCCTTTCTCCCTTCGGGAAGCGTGGCGCTTTCTCAATGCTCACGCTGTAGGTATCTCAGTTCGGTGTAGGTCGTTCGCTCCAAGCTGGGCTGTGTGCACGAACCCCCCGTTCAGCCCGACCGCTGCGCCTTATCCGGTAACTATCGTCTTGAGTCCAACCCGGTAAGACACGACTTATCGCCACTGGCAGCAGCCACTGGTAACAGGATTAGCAGAGCGAGGTATGTAGGCGGTGCTACAGAGTTCTTGAAGTGGTGGCCTAACTACGGCTACACTAGAAGGACAGTATTTGGTATCTGCGCTCTGCTGAAGCCAGTTACCTTCGGAAAAAGAGTTGGTAGCTCTTGATCCGGCAAACAAACCACCGCTGGTAGCGGTGGTTTTTTTGTTTGCAAGCAGCAGATTACGCGCAGAAAAAAAGGATCTCAAGAAGATCCTTTGATCTTTTCTACGGGGTCTGACGCTCAGTGGAACGAAAACTCACGTTAAGGGATTTTGGTCATGAGATTATCAAAAAGGATCTTCACCTAGATCCTTTTAAATTAAAAATGAAGTTTTAAATCAATCTAAAGTATATATGAGTAAACTTGGTCTGACAGTAATGCCCTGCACTTCATCCTTAACTGGTATCGGGACTCTTAGGTGAGTGGTGAGATACGGGGCATGATGTCCCCTTCCCCCTTTGGGTTTATCTATGCCCTGCATGGGTTTAGCCTGTTAACGGGATTATATTCCCATCATAACTCCCTGTTTTTGGTATCCAGTTCCTAGGCTTGATTAATTAATAAGGATTCAGTGGATACGGTATCAGAGTGATACAAAATAGAATCCCGCTCAAACCACCGGGAAAAAATAGCCGATCGCCCATCGGGGTCAATATTGGCGGGGCCATAATGTCGCTTAGTATTGCCCCGTTCCCCAAACCGGCCCAGATACTCAAGATGTAAACCCATTTTTTTCAAGACAGATTGCACGAAGGCGATCGCCCCTTTGTCAGGGTTGATCGTGATGCCCAAGGCCTGCTTAATTTCTGCCTTGTTGGGCAGGATAGTCTTCTCAAACCAATTGGCTAGGGAATCCTTGGAAAATTCCCCTATCCCGGTAAGGAATCTTTCAAAGCCCAATACTTTAAGGAAGTGAACGGGGACGGTGAGAGTCTTCCGGCTAGCATCCTTGGCACAAACCTTTCCATCTTCCCCGGCCAAATTTTGGAGCCTTTGGCTGTCCCTTGCCTGTAAAAATTCTGCACCGGTGGTGAAGTAATAATGCAGGGTCAATTGGGCCAGCCATCCTTGATCACATTTCTCCACATCTTCTGGGGTTATGTCATCGGTAGCCAGGGCACGGCAAAGATCTCCCTTCCGCTCTTGATATCGTTCCGCTTCCGTCTTAACTTGCTTCCGCTTCAAACTTTCATAGGTGAAGTCATCGGGATTAAGGCTCTCGGCGATCGCCGTGGTTTCCTTTCCATAGTTATGGTCACGGATAGCTTTAATGCGATCAGCTACGTCTTCCGCCCCGGTGACCATATCTGGGTCAGGTTCTTTCTGTGTGTACCCTTCTTCCCCTAATTTTTCAAGAGTAGTTGATTTTAAGCTTTTCTTCCCTTGATTGTGCAGAGCCGCCATAATAGCCCAGGTTTTAAGATGCTCTGGCTTCTCATCATCAAACAGGGTTAAATTGTCGGCTGTATTCAATGCCCCGATAGTAGCCTTGGCTTTCTTATTCTCCACATTAAGGATATAGCGAGGGTTAGTTTCCCCGCCAGCAATAAAACAATTATTAGGGGCACGATTGCGAATCCAAACATGGCGATCCACATCGGGACGATACCTTTCTAATCCTTGGCAGAATTGCTCTACTGTTTGTGTTCCCTGTCCGAATCCATAGACTTCATCAAAATAGGGCTTATCAATGCTGATTCCCGTCTCTATTACCGGGGACGTGATGACAATATCCTTGTCTTCTAAATATCTATCTAAGTTATCCATACAGCCATAGGCAGGGTTAGTCTTATCGCTTACTGTATGGGCATCTAAGACCCCTACTGACTTATCAGGGAATAACATTTTGAATAAGTCCCCTAGGTTAGTTGTTGAATAGATAGACTGTGCTTTTTGTGCCCCGGTGCAAATCTGTATTTTTTTACCATTAGCAACAGTATTAATAGCGGAAGTAAGCAAATCTTCCGGGGAATTATAAACAATCAAATTACGCTTATCCTGCACTGGCTTAAACGTGTTAACGACAATATAGCATTCTGCCTTACCACCGATTAAGTCTTGGACATACTTGATAGTCACCGGGGACAAATCAGCATCAGACAAAATAATCTTGCCCCCCGAATCAGCGGCGGCGATCAATATTTCTTGGAAGCGGGCCAAGATGGCTGGTTTATGTTTGCCCAGGTTTCCCCGGGAACTATCTAATAATTCCCATATCAATTGCTCGCATTCATCAAGGATGATGTCATGGCCCCGGAATGTTTCGGGGTCAAGTTTCAGGATGCTATCAATGCACAATCCGATTCCTAAGGCCCCATTAGTGTCGCTCTCGGACAAATCTCTGATGTAGTCCACACCAAACCGGCCGGATAGGGCAATGCCTAGTTGAATTCGGTGAGTGAGTGGATAAGTCCGGCGAAAATCTTCTCGCCGTTGCTCCACCATTTTTGAGATCGCTTCCGTCTTTCCCGTGCCCTTGGCTGATTTAATCCCGATTATTTTGGCGGTGGTAGGGGCATTAATGTTGCTTAGGTATCTTTCGTGCCGGGTTTGATTCACCCATTGGGATAGGTCAGCATACGGCTTGTTTTGGAACTGTTCTAGGCTGATGCCCCGAGCGATCGCCCGGTGAAATTTAGCAGGGTTATTGGCGATCAAGTCATCAATGCCCTTGCCGTCTTCCCCGTTCCATTGGCAGATCTTGACCGTTGCCTTAGCGTGATAGGTCAGGTTACGAGCCAGCCGTTTTGTGCCCTTAAAGACTGCCTTTCGTCCCTTGCTACCCTTTGCATCCTGGTCATAGGCGATCGCTACTTCTCGCCCCTGCACATAGGGCAATAATGACGGTTTGATGGTTAGACCATCGTTGCCACATAAACAGCCATAGAGACTCAGGGCCACGTAACCTTGGCTGATGGCGGCTAAAGCTTTCTTCCCCCCTTCCGTCACAATCAAGGGAATCTCAGGATGATCCACAAACCAGGGCCAAAAATCTTGACCATCTTCCGGTGGTTTGACCCCATGCTTTTCGGCGATCGCCAAAATGATCCGGCGGGGAATTGTTGGCAGATAGGGCACGTCCCCTATGCCTGTCGGTGCTAGATACTGCCCTGACCGCTTGCCCGTCTTGTCTTCACCAAAGATTTTTACCTGCCATACGTTCCCATTCTCAGACCGGAAAATGGCGGCCTGCCTGTCTTCCTTGGCTTGATGTCCAAAGCGGGTAAATTTTCGCCCCAGGGCATCGGCGATCGGCGTTGCCAAGACTTCCTTGGTGATGGGGTCTATCTCAAGATCCGGCACAATCTCCACGTTGGCCCGGAATACGTCAGGGGCGATCGCCGAATCTTCTACAAATTCTGACCGTATTTTTGGGTCATTTTTTGACCCTTGCCATCCGTTGAAATGCTGATGGGCTAAGGCTTGAGAATTTTGGTGGGAAAGACTATAATTGTTCATAGCGGTCTCGGGTTCAAGAGTTGACCGCTTTTCTTTTGGGAATTTTTAGAGCTACAGTAGAGACAATATTTTTGAAGTATTTTGTTGGCTTTTTGGGCCGGGTTATTGTCTGTAGGTAAGTTTTTTAGCGTCAACTAAACTTCTTACGTTTTTTACTGCAGCAATCTTAATTCCCTCCTGAAAAGCTTTTTTTTGTTGGAAATCTTGGACTATTGCCAGAATTGTAACGGATTAGATCGGTCGCTGTCACGCTGTTTTATCGTCATAAAAGTTTATTTAAATTAAATATGTGAGCAATGCTCACGGCTGTCGCTATCCCTTTGTCAAGAGATAATACACTTTGGATTGTCCCCTAAAAGATTGTTTTTATAATCGGCTTATTTTCTCTTTCTGGTGTTATTAGTCTCTTTTTGAGCGACGACGATC

Complete nucleotide sequence of pYS1C-GFP. *trc* promoter (red), lacO (gray), *gfp*^mut2^ (green), *cat* gene (pink), *ColE* (light blue), pCC5.2 region (yellow)

**Supplementary material Data 2**

CCAGGCATCAAATAAAACGAAAGGCTCAGTCGAAAGACTGGGCCTTTCGTTTTATCTGTTGTTTGTCGGTGAACGCTCTCTACTAGAGTCACACTGGCTCACCTTCGGGTGGGCCTTTCTGCGTTTATACATTAATTGCGTTGCGCTCACTGCCCGCTTTCCAGTCGGGAAACCTGTCGTGCCAGCTGCATTAATGAATCGGCCAACGCGCGGGGAGAGGCGGTTTGCGTATTGGGCGCCAGGGTGGTTTTTCTTTTCACCAGTGAGACGGGCAACAGCTGATTGCCCTTCACCGCCTGGCCCTGAGAGAGTTGCAGCAAGCGGTCCACGCTGGTTTGCCCCAGCAGGCGAAAATCCTGTTTGATGGTGGTTAACGGCGGGATATAACATGAGCTGTCTTCGGTATCGTCGTATCCCACTACCGAGATATCCGCACCAACGCGCAGCCCGGACTCGGTAATGGCGCGCATTGCGCCCAGCGCCATCTGATCGTTGGCAACCAGCATCGCAGTGGGAACGATGCCCTCATTCAGCATTTGCATGGTTTGTTGAAAACCGGACATGGCACTCCAGTCGCCTTCCCGTTCCGCTATCGGCTGAATTTGATTGCGAGTGAGATATTTATGCCAGCCAGCCAGACGCAGACGCGCCGAGACAGAACTTAATGGGCCCGCTAACAGCGCGATTTGCTGGTGACCCAATGCGACCAGATGCTCCACGCCCAGTCGCGTACCGTCTTCATGGGAGAAAATAATACTGTTGATGGGTGTCTGGTCAGAGACATCAAGAAATAACGCCGGAACATTAGTGCAGGCAGCTTCCACAGCAATGGCATCCTGGTCATCCAGCGGATAGTTAATGATCAGCCCACTGACGCGTTGCGCGAGAAGATTGTGCACCGCCGCTTTACAGGCTTCGACGCCGCTTCGTTCTACCATCGACACCACCACGCTGGCACCCAGTTGATCGGCGCGAGATTTAATCGCCGCGACAATTTGCGACGGCGCGTGCAGGGCCAGACTGGAGGTGGCAACGCCAATCAGCAACGACTGTTTGCCCGCCAGTTGTTGTGCCACGCGGTTGGGAATGTAATTCAGCTCCGCCATCGCCGCTTCCACTTTTTCCCGCGTTTTCGCAGAAACGTGGCTGGCCTGGTTCACCACGCGGGAAACGGTCTGATAAGAGACACCGGCATACTCTGCGACATCGTATAACGTTACTGGTTTCACATTCACCACCCTGAATTGACTCTCTTCCGGGCGCTATCATGCCATACCGCGAAAGGTTTTGCGCCATTCGATGGTGTCCGGGATCTCGACGCTCTCCCTTATGCGACTCCTGCATTAGGAAGCAGCCCAGTAGTAGGTTGAGGCCGTTGAGCACCGCCGCCGCAAGGAATGGTGCATGCAAGGAGATGGCGCCCAACAGTCCCCCGGCCACGGGGCCTGCCACCATACCCACGCCGAAACAAGCGCTCATGAGCCCGAATTGTGAGCGCTCACAATTCGGAATTCTTAACAAAAAAGCAGGAATAAAATTAACAAGATGTAATTGACATAAGTCCCATCACCGTTGTATAAATGTGTGGAATTGTGAGCGGATAACAATTTCACACAATGGAATTCAGTAAAGGAGAAGAACTTTTCACTGGAGTTGTCCCAATTCTTGTTGAATTAGATGGTGATGTTAATGGGCACAAATTTTCTGTCAGTGGAGAGGGTGAAGGTGATGCAACATACGGAAAACTTACCCTTAAATTTATTTGCACTACTGGAAAACTACCTGTTCCATGGCCAACACTTGTCACTACTTTCGCGTATGGTCTTCAATGCTTTGCGAGATACCCAGATCATATGAAACAGCATGACTTTTTCAAGAGTGCCATGCCCGAAGGTTATGTACAGGAAAGAACTATATTTTTCAAAGATGACGGGAACTACAAGACACGTGCTGAAGTCAAGTTTGAAGGTGATACCCTTGTTAATAGAATCGAGTTAAAAGGTATTGATTTTAAAGAAGATGGAAACATTCTTGGACACAAATTGGAATACAACTATAACTCACACAATGTATACATCATGGCAGACAAACAAAAGAATGGAATCAAAGTTAACTTCAAAATTAGACACAACATTGAAGATGGAAGCGTTCAACTAGCAGACCATTATCAACAAAATACTCCAATTGGCGATGGCCCTGTCCTTTTACCAGACAACCATTACCTGTCCACACAATCTGCCCTTTCGAAAGATCCCAACGAAAAGAGAGACCACATGGTCCTTCTTGAGTTTGTAACAGCTGCTGGGATTACACATGGCATGGATGAACTATACAAATAAGCTTACTAGTAATACTGCAGAGAGAATATAAAAAGCCAGATTATTAATCCGGCTTTTTTATTATTTAGACGTCAGGTGGCACTTTTCGGGGAAATGTGAGAATAAATAAATCCTGGTGTCCCTGTTGATACCGGGAAGCCCTGGGCCAACTTTTGGCGAAAATGAGACGTTGATCGGCACGTAAGAGGTTCCAACTTTCACCATAATGAAATAAGATCACTACCGGGCGTATTTTTTGAGTTATCGAGATTTTCAGGAGCTAAGGAAGCTAAAATGGAGAAAAAAATCACTGGATATACCACCGTTGATATATCCCAATGGCATCGTAAAGAACATTTTGAGGCATTTCAGTCAGTTGCTCAATGTACCTATAACCAGACCGTTCAGCTGGATATTACGGCCTTTTTAAAGACCGTAAAGAAAAATAAGCACAAGTTTTATCCGGCCTTTATTCACATTCTTGCCCGCCTGATGAATGCTCATCCGGAATTCCGTATGGCAATGAAAGACGGTGAGCTGGTGATATGGGATAGTGTTCACCCTTGTTACACCGTTTTCCATGAGCAAACTGAAACGTTTTCATCGCTCTGGAGTGAATACCACGACGATTTCCGGCAGTTTCTACACATATATTCGCAAGATGTGGCGTGTTACGGTGAAAACCTGGCCTATTTCCCTAAAGGGTTTATTGAGAATATGTTTTTCGTCTCAGCCAATCCCTGGGTGAGTTTCACCAGTTTTGATTTAAACGTGGCCAATATGGACAACTTCTTCGCCCCCGTTTTCACCATGGGCAAATATTATACGCAAGGCGACAAGGTGCTGATGCCGCTGGCGATTCAGGTTCATCATGCCGTTTGTGATGGCTTCCATGTCGGCAGAATGCTTAATGAATTACAACAGTACTGCGATGAGTGGCAGGGCGGGGCGTAACAACATACGAGCCGGAAGCATAAAGTGTAAAGCCTGGGGTGCCTAATGAGTGAGCTAACTCACATTAATTGCGTTGCGCTCACTGCCCGCTTTCCAGTCGGGAAACCTGTCGTGCCAGCTGCATTAATGAATCGGCCAACGCGCGGGGAGAGGCGGTTTGCGTATTGGGCGCTCTTCCGCTTCCTCGCTCACTGACTCGCTGCGCTCGGTCGTTCGGCTGCGGCGAGCGGTATCAGCTCACTCAAAGGCGGTAATACGGTTATCCACAGAATCAGGGGATAACGCAGGAAAGAACATGTGAGCAAAAGGCCAGCAAAAGGCCAGGAACCGTAAAAAGGCCGCGTTGCTGGCGTTTTTCCATAGGCTCCGCCCCCCTGACGAGCATCACAAAAATCGACGCTCAAGTCAGAGGTGGCGAAACCCGACAGGACTATAAAGATACCAGGCGTTTCCCCCTGGAAGCTCCCTCGTGCGCTCTCCTGTTCCGACCCTGCCGCTTACCGGATACCTGTCCGCCTTTCTCCCTTCGGGAAGCGTGGCGCTTTCTCAATGCTCACGCTGTAGGTATCTCAGTTCGGTGTAGGTCGTTCGCTCCAAGCTGGGCTGTGTGCACGAACCCCCCGTTCAGCCCGACCGCTGCGCCTTATCCGGTAACTATCGTCTTGAGTCCAACCCGGTAAGACACGACTTATCGCCACTGGCAGCAGCCACTGGTAACAGGATTAGCAGAGCGAGGTATGTAGGCGGTGCTACAGAGTTCTTGAAGTGGTGGCCTAACTACGGCTACACTAGAAGGACAGTATTTGGTATCTGCGCTCTGCTGAAGCCAGTTACCTTCGGAAAAAGAGTTGGTAGCTCTTGATCCGGCAAACAAACCACCGCTGGTAGCGGTGGTTTTTTTGTTTGCAAGCAGCAGATTACGCGCAGAAAAAAAGGATCTCAAGAAGATCCTTTGATCTTTTCTACGGGGTCTGACGCTCAGTGGAACGAAAACTCACGTTAAGGGATTTTGGTCATGAGATTATCAAAAAGGATCTTCACCTAGATCCTTTTAAATTAAAAATGAAGTTTTAAATCAATCTAAAGTATATATGAGTAAACTTGGTCTGACAGTAATGCCCTGCACTTCATCCTTAACTGGTATCGGGACTCTTAGGTGAGTGGTGAGATACGGGGCATGATGTCCCCTTCCCCCTTTGGGTTTATCTATGCCCTGCATGGGTTTAGCCTGTTAACGGGATTATATTCCCATCATAACTCCCTGTTTTTGGTATCCAGTTCCTAGGCTTGATTAATTAATAAGGATTCAGTGGATACGGTATCAGAGTGATACAAAATAGAATCCCGCTCAAACCACCGGGAAAAAATAGCCGATCGCCCATCGGGGTCAATATTGGCGGGGCCATAATGTCGCTTAGTATTGCCCCGTTCCCCAAACCGGCCCAGATACTCAAGATGTAAACCCATTTTTTTCAAGACAGATTGCACGAAGGCGATCGCCCCTTTGTCAGGGTTGATCGTGATGCCCAAGGCCTGCTTAATTTCTGCCTTGTTGGGCAGGATAGTCTTCTCAAACCAATTGGCTAGGGAATCCTTGGAAAATTCCCCTATCCCGGTAAGGAATCTTTCAAAGCCCAATACTTTAAGGAAGTGAACGGGGACGGTGAGAGTCTTCCGGCTAGCATCCTTGGCACAAACCTTTCCATCTTCCCCGGCCAAATTTTGGAGCCTTTGGCTGTCCCTTGCCTGTAAAAATTCTGCACCGGTGGTGAAGTAATAATGCAGGGTCAATTGGGCCAGCCATCCTTGATCACATTTCTCCACATCTTCTGGGGTTATGTCATCGGTAGCCAGGGCACGGCAAAGATCTCCCTTCCGCTCTTGATATCGTTCCGCTTCCGTCTTAACTTGCTTCCGCTTCAAACTTTCATAGGTGAAGTCATCGGGATTAAGGCTCTCGGCGATCGCCGTGGTTTCCTTTCCATAGTTATGGTCACGGATAGCTTTAATGCGATCAGCTACGTCTTCCGCCCCGGTGACCATATCTGGGTCAGGTTCTTTCTGTGTGTACCCTTCTTCCCCTAATTTTTCAAGAGTAGTTGATTTTAAGCTTTTCTTCCCTTGATTGTGCAGAGCCGCCATAATAGCCCAGGTTTTAAGATGCTCTGGCTTCTCATCATCAAACAGGGTTAAATTGTCGGCTGTATTCAATGCCCCGATAGTAGCCTTGGCTTTCTTATTCTCCACATTAAGGATATAGCGAGGGTTAGTTTCCCCGCCAGCAATAAAACAATTATTAGGGGCACGATTGCGAATCCAAACATGGCGATCCACATCGGGACGATACCTTTCTAATCCTTGGCAGAATTGCTCTACTGTTTGTGTTCCCTGTCCGAATCCATAGACTTCATCAAAATAGGGCTTATCAATGCTGATTCCCGTCTCTATTACCGGGGACGTGATGACAATATCCTTGTCTTCTAAATATCTATCTAAGTTATCCATACAGCCATAGGCAGGGTTAGTCTTATCGCTTACTGTATGGGCATCTAAGACCCCTACTGACTTATCAGGGAATAACATTTTGAATAAGTCCCCTAGGTTAGTTGTTGAATAGATAGACTGTGCTTTTTGTGCCCCGGTGCAAATCTGTATTTTTTTACCATTAGCAACAGTATTAATAGCGGAAGTAAGCAAATCTTCCGGGGAATTATAAACAATCAAATTACGCTTATCCTGCACTGGCTTAAACGTGTTAACGACAATATAGCATTCTGCCTTACCACCGATTAAGTCTTGGACATACTTGATAGTCACCGGGGACAAATCAGCATCAGACAAAATAATCTTGCCCCCCGAATCAGCGGCGGCGATCAATATTTCTTGGAAGCGGGCCAAGATGGCTGGTTTATGTTTGCCCAGGTTTCCCCGGGAACTATCTAATAATTCCCATATCAATTGCTCGCATTCATCAAGGATGATGTCATGGCCCCGGAATGTTTCGGGGTCAAGTTTCAGGATGCTATCAATGCACAATCCGATTCCTAAGGCCCCATTAGTGTCGCTCTCGGACAAATCTCTGATGTAGTCCACACCAAACCGGCCGGATAGGGCAATGCCTAGTTGAATTCGGTGAGTGAGTGGATAAGTCCGGCGAAAATCTTCTCGCCGTTGCTCCACCATTTTTGAGATCGCTTCCGTCTTTCCCGTGCCCTTGGCTGATTTAATCCCGATTATTTTGGCGGTGGTAGGGGCATTAATGTTGCTTAGGTATCTTTCGTGCCGGGTTTGATTCACCCATTGGGATAGGTCAGCATACGGCTTGTTTTGGAACTGTTCTAGGCTGATGCCCCGAGCGATCGCCCGGTGAAATTTAGCAGGGTTATTGGCGATCAAGTCATCAATGCCCTTGCCGTCTTCCCCGTTCCATTGGCAGATCTTGACCGTTGCCTTAGCGTGATAGGTCAGGTTACGAGCCAGCCGTTTTGTGCCCTTAAAGACTGCCTTTCGTCCCTTGCTACCCTTTGCATCCTGGTCATAGGCGATCGCTACTTCTCGCCCCTGCACATAGGGCAATAATGACGGTTTGATGGTTAGACCATCGTTGCCACATAAACAGCCATAGAGACTCAGGGCCACGTAACCTTGGCTGATGGCGGCTAAAGCTTTCTTCCCCCCTTCCGTCACAATCAAGGGAATCTCAGGATGATCCACAAACCAGGGCCAAAAATCTTGACCATCTTCCGGTGGTTTGACCCCATGCTTTTCGGCGATCGCCAAAATGATCCGGCGGGGAATTGTTGGCAGATAGGGCACGTCCCCTATGCCTGTCGGTGCTAGATACTGCCCTGACCGCTTGCCCGTCTTGTCTTCACCAAAGATTTTTACCTGCCATACGTTCCCATTCTCAGACCGGAAAATGGCGGCCTGCCTGTCTTCCTTGGCTTGATGTCCAAAGCGGGTAAATTTTCGCCCCAGGGCATCGGCGATCGGCGTTGCCAAGACTTCCTTGGTGATGGGGTCTATCTCAAGATCCGGCACAATCTCCACGTTGGCCCGGAATACGTCAGGGGCGATCGCCGAATCTTCTACAAATTCTGACCGTATTTTTGGGTCATTTTTTGACCCTTGCCATCCGTTGAAATGCTGATGGGCTAAGGCTTGAGAATTTTGGTGGGAAAGACTATAATTGTTCATAGCGGTCTCGGGTTCAAGAGTTGACCGCTTTTCTTTTGGGAATTTTTAGAGCTACAGTAGAGACAATATTTTTGAAGTATTTTGTTGGCTTTTTGGGCCGGGTTATTGTCTGTAGGTAAGTTTTTTAGCGTCAACTAAACTTCTTACGTTTTTTACTGCAGCAATCTTAATTCCCTCCTGAAAAGCTTTTTTTTGTTGGAAATCTTGGACTATTGCCAGAATTGTAACGGATTAGATCGGTCGCTGTCACGCTGTTTTATCGTCATAAAAGTTTATTTAAATTAAATATGTGAGCAATGCTCACGGCTGTCGCTATCCCTTTGTCAAGAGATAATACACTTTGGATTGTCCCCTAAAAGATTGTTTTTATAATCGGCTTATTTTCTCTTTCTGGTGTTATTAGTCTCTTTTTGAGCGACGACGATC

Complete nucleotide sequence of pYS4C-GFP. *cLac143* promoter (red), lacO (gray), *gfp*^mut2^ (green), *cat* gene (pink), *ColE* (light blue), pCC5.2 region (yellow)
